# Differential TIM-3 glycosylation enables specific dual targeting CAR-T therapy in acute myeloid leukemia

**DOI:** 10.64898/2026.04.22.719217

**Authors:** Marta Biondi, Antonio Galeone, Corinne Arsuffi, Bruna My, Erica Grassenis, Camilla Firpo, Maria Caterina Rotiroti, Alice Pievani, Antonio Maria Alviano, Beatrice Cerina, Giuseppe Gigli, Andrea Lia, Andrea Biondi, Sarah Tettamanti, Marta Serafini

## Abstract

Chimeric Antigen Receptor (CAR) therapy for Acute Myeloid Leukemia (AML) is hindered by disease heterogeneity, antigen overlap with normal hematopoiesis, and the persistence of leukemic stem cells (LSCs). To overcome these barriers, we employed the IF-BETTER strategy to simultaneously target CD33 and the LSC-associated marker TIM-3 by pairing a second-generation CAR with a Cytokine-Costimulatory Receptor (CCR) in two dual CAR configurations (CD33.CAR/TIM-3.CCR and TIM-3.CAR/CD33.CCR). Both constructs displayed potent antigen-restricted cytotoxicity against AML cell lines and primary blasts, achieving leukemia clearance, while sparing normal immune and hematopoietic cells. Mechanistic studies revealed that the TIM-3.CAR single-chain fragment variable (scFv) recognizes a protein-proximal epitope whose interaction is selectively enhanced by AML-specific hyper-fucosylated and hyper-sialylated N-glycans. Fucosylation blockade reduced TIM-3.CAR avidity and cytotoxicity, confirming a glycosylation-modulated interaction. Integrating this glycosylation-tolerant TIM-3 scFv into a dual CAR framework enables selective targeting of AML cells, providing a rational strategy for safer and more effective AML-directed immunotherapy.

## INTRODUCTION

Despite significant advances in molecular stratification and supportive care^1–3^, Acute Myeloid Leukemia (AML) is still associated with high rates of relapse and refractory disease, largely driven by therapy-resistant leukemic stem cells (LSCs) that persist within the bone marrow (BM) niche^4^. As a result, 5-year overall survival remains ∼50% in younger adults and drops below 10% in patients over 60 years of age^3,5^. Allogenic hematopoietic stem cell transplantation (allo-HSCT) is the only approved curative treatment, yet many patients are ineligible or experience severe toxicities, and relapse after transplant remains frequent^6^. Thus, more effective and durable therapies for AML are urgently needed.

Chimeric Antigen Receptor (CAR)-T cell therapy offers a promising opportunity to redirect endogenous immune responses directly against AML blasts. However, despite their remarkable success in B-cell malignancies^7,8^, CAR-T cells have shown limited clinical efficacy in AML^9^. Major hurdles include the scarcity of LSC/AML-specific target antigens, the disease heterogeneity and the leukemia-mediated remodeling of the BM microenvironment^10^. Early clinical trials targeting overexpressed shared antigens, such as CD33 or CD123, have been constrained by insufficient specificity, poor persistence, limited therapeutic benefit and life-threatening toxicities^11,12^.

These limitations have prompted a shift toward identifying more LSC-restricted antigens and developing next-generation CAR designs^13^. T-cell Immunoglobulin Mucin-3 (TIM-3) has emerged as a promising LSC-selective target^14–17:^ its expression correlates with poor prognosis^18^, it is absent on healthy hematopoietic stem cells (HSCs)^19^ and its engagement in an autocrine TIM-3/Galectin-9 (Gal-9) loop enhances LSC stemness, self-renewal, and immune evasion^20–22^. However, its constitutive expression on multiple innate and adaptive immune cells, such as monocytes and activated T and NK cells^17^, raises concerns for on-target/off-tumor toxicity.

Here, we develop and characterize a novel TIM-3.CAR derived from an antagonistic ligand-blocking monoclonal antibody recognizing a protein-proximal TIM-3 epitope whose binding, as we demonstrate, is selectively potentiated by AML-associated *N*-glycan modifications. We show that AML cells display hyper-fucosylated and hyper-sialylated TIM-3 glycoforms that are absent on healthy TIM-3^+^ immune cells, and that blockade of fucosylation reduces TIM-3.CAR binding and cytotoxicity. Notably, these aberrant glycosylation patterns are widely recognized hallmarks of solid and hematological tumors, including AML, playing a crucial role in cancer initiation and progression by promoting tumor evasion, metastasis, and chemoresistance^23–26^. These findings reveal that AML-specific glycosylation functionally enhances CAR-antigen avidity, providing a structural and biochemical rationale for the selective engagement of leukemic TIM-3 and sparing of healthy tissues.

To translate this selectivity into a therapeutic strategy suited to the intrinsic challenges of AML, we selected TIM-3 as an LSC-enriched antigen and paired it with CD33, a well-established and broadly expressed AML marker^27,28^. We incorporated the glycosylation-tolerant TIM-3 single-chain fragment variable (scFv) into an IF-BETTER combinatorial logic framework by pairing a second-generation CAR with a cytokine-costimulatory receptor (CCR). This antigen combination was designed to exploit TIM-3 LSC specificity while leveraging CD33 robust coverage of the bulk leukemic compartment. Dual CAR configurations co-targeting CD33 and TIM-3 (CD33.CAR/TIM-3.CCR and TIM-3.CAR/CD33.CCR) demonstrated potent and selective cytotoxicity against AML cell lines and patient-derived samples, mediated efficient leukemia control *in vivo*, and spared healthy hematopoietic and immune cells. This combinatorial approach increases the precision of AML targeting, mitigates on-target/off-tumor toxicity, and contributes to overcoming intratumoral heterogeneity.

Together, our findings suggest that integrating a glycosylation-tolerant scFv, whose binding is selectively enhanced by AML-specific *N*-glycan modifications, into an IF-BETTER dual CAR platform represents a promising therapeutic strategy for AML.

## RESULTS

### TIM-3 is selectively expressed on primary AML blasts and LSC-enriched compartments

To validate TIM-3 as a selective AML target, we first analyzed its expression in primary samples from 25 AML patients. The cohort included both adult and pediatric patients representing different molecular and genetic subtypes. In accordance with previous reports^14,15,19^, TIM-3 was consistently enriched on AML blasts as compared to residual healthy CD45^high^ bone marrow cells (**Figure 1A**). Moreover, a substantial fraction of LSC-enriched population (CD34^+^CD38^-^) also expressed TIM-3 (**Figure 1B**). These findings support TIM-3 as a discriminatory marker across AML differentiation states.

**Figure 1.**
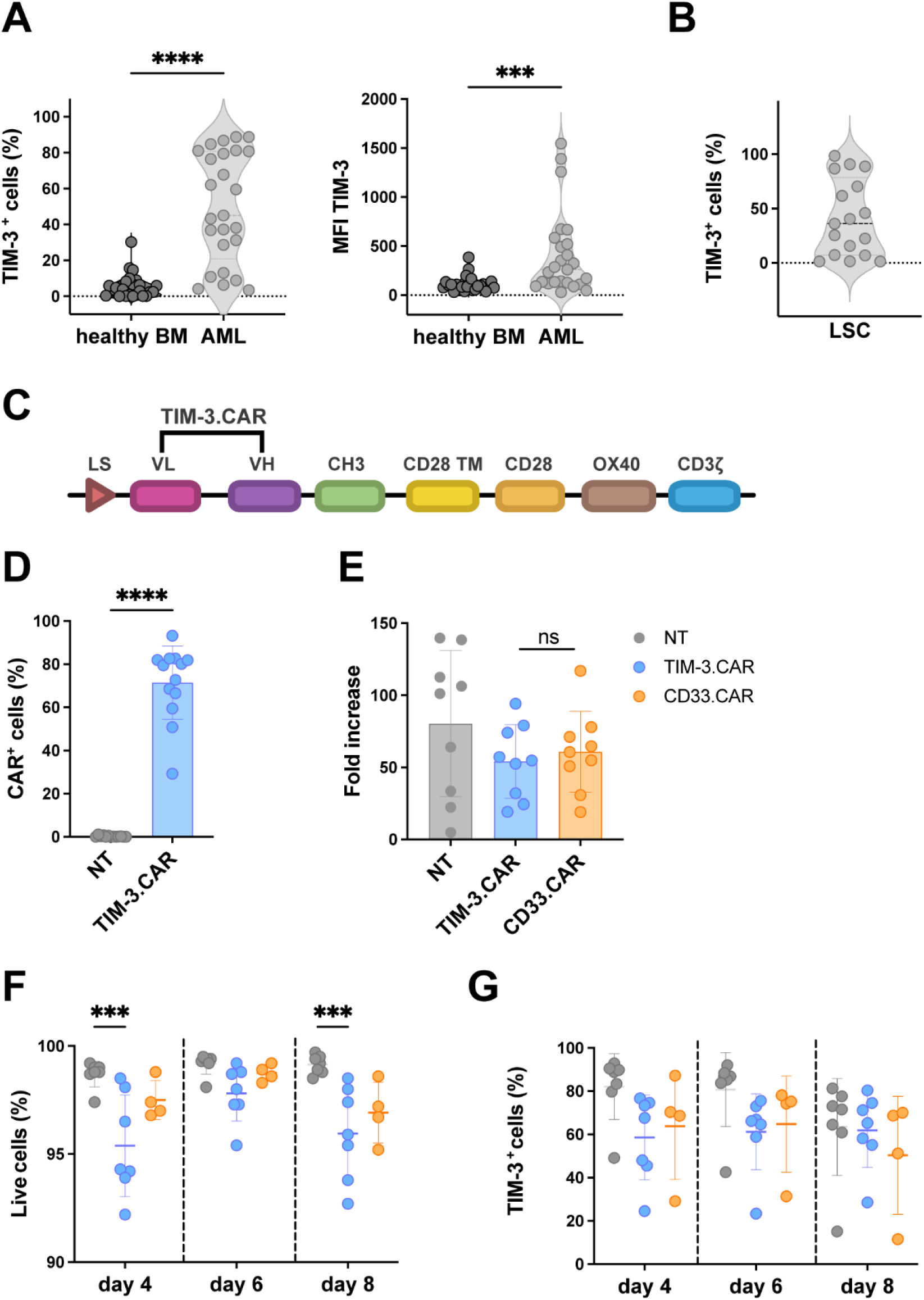
TIM-3.CAR is efficiently expressed in CIK cells without inducing fratricide. **(A)** Frequency (left) and mean fluorescence intensity (MFI; right) of TIM-3 expression on bulk AML blasts compared with residual healthy bone marrow (BM) cells from the same patient (n=25). **(B)** TIM-3 expression in the LSC-enriched CD34^+^ CD38^-^ population (n=17). **(C)** Schematic of the third-generation TIM-3.CAR containing CD28-OX40 costimulatory domains cloned into the pT4-transposon vector. **(D)** Transduction efficiency of TIM-3.CAR-CIK cells compared to non-transduced (NT) CIK cells (mean 71.24±16.99, n = 13). See also **Figure S1A**. **(E)** Fold increase of TIM-3.CAR-CIK cells compared to CD33.CAR-CIK and NT cells at day 21 of culture (n=9). **(F)** Viability of TIM-3.CAR-CIK cells, CD33.CAR-CIK and NT cells at days 4, 6 and 8 post-transduction, assessed by flow cytometry. See also **Figure S1B**. **(G)** Frequency of CD3^+^TIM-3^+^ cells within NT, TIM3.CAR- and CD33.CAR-CIK populations over the first 8 days of CIK differentiation (n=7 donors for TIM-3.CAR-CIK and NT cells, n=4 donors for CD33.CAR-CIK cells). See also **Figure S1C**. Data are shown as individual values with mean ± standard deviation (SD). Significance was assessed using paired t test (A, D) or repeated-measures two-way ANOVA with Bonferroni’s post hoc test (E-G). ns, not significant, *** p = 0.0001 and **** p < 0.0001. See also **Figure S1** for longitudinal TIM-3 and TIM-3.CAR expression and CD4^+^ and CD8^+^ subset analysis.

### Efficient expansion of TIM-3.CAR-CIK cells despite endogenous TIM-3 expression

We generated a third-generation anti-TIM-3.CAR comprising the scFv of an antagonistic ligand-blocking anti-TIM-3 antibody^29^ (clone M6903) cloned in frame with the endodomain of the CAR carrying the CD28-OX40 costimuli within the Sleeping Beauty (SB) pT4-transposon plasmid (**Figure 1C**). TIM-3.CAR cytokine-induced-killer (CIK) cells, which are *ex vivo* expanded T lymphocytes with a mixed T–natural killer (NK) phenotype endowed with MHC-unrestricted antitumor activity, were successfully generated across donors following a validated protocol^30^, achieving high and stable CAR expression (up to 70%) during expansion (n=13 donors) (**Figure 1D**).

Considering the physiological expression of TIM-3 on activated T cells^17^, we evaluated the potential fratricide effect of TIM-3.CAR-CIK cells. TIM-3.CAR-CIK cells expanded efficiently, comparably to CD33.CAR-CIK controls (**Figure 1E**) and maintained normal viability, with no impairment in proliferative kinetics, suggesting the absence of an overt fratricide (**Figure 1F**). Notably, TIM-3 expression was detectable from the initial days of culture, suggesting that a TIM-3-negative population was not selectively enriched (**Figure 1G**). To further investigate this potential issue, we followed TIM-3.CAR expression together with TIM-3 expression during the first 8 days of CAR-CIK differentiation. TIM-3 levels remained stable across CD4⁺ and CD8⁺ subsets despite a modest shift toward lower TIM-3 expression levels, while TIM-3.CAR showed a progressive enrichment during culture. Cell viability remained consistently high, with minor variations likely attributable to the recent manipulation rather than to active fratricide (**Figures S1A–1C**). Collectively, these findings demonstrate robust expansion of TIM-3.CAR-CIK cells despite endogenous TIM-3 expression.

### TIM-3.CAR-CIK cells selectively and potently eliminate TIM-3⁺ AML cells in vitro

To assess the antileukemic activity of TIM-3.CAR-CIK cells, we selected the TIM-3^+^ AML cell line KASUMI-3 and adopted the TIM-3^-^ acute lymphoblastic leukemia (ALL) cell line REH as a negative control (**Figure S2A**). TIM-3.CAR-CIK cells mediated strong and antigen-specific cytotoxicity in short-term assays, with efficient clearance of TIM-3⁺ KASUMI-3 target and minimal activity against TIM-3⁻ REH cells (**Figures S2B–2C**). In long-term co-culture, TIM-3.CAR-CIK cells sustained potent TIM-3–dependent killing across limiting effector-to-target (E:T) ratios and expanded with a superior proliferation index as compared to controls in response to leukemic stimulation without exogenous IL-2 cytokine support (**Figures S2D–2F**). Furthermore, TIM-3.CAR engagement also induced antigen-driven proliferation and robust IFN-γ and IL-2 production, whereas stimulation with TIM-3⁻ targets elicited no appreciable responses (**Figures S2G–2H**).

TIM-3 and PD-1 are well-established inhibitory receptors associated with T cell exhaustion^31^ and CAR-T cells dysfunctions in AML^32^. In particular, AML TIM-3/Gal-9 circuits can engage TIM-3 on T cells and reinforce these exhaustion pathways^20–22^. Notably, KASUMI-3 cells and primary AML blasts express elevated levels of Gal-9 compared to the control REH cell line (**Figure S2I**). We therefore investigated whether targeting TIM-3 through the CAR could modulate this AML-driven exhaustion loop, by analyzing TIM-3 and PD-1 expression before and after long-term co-culture experiments. TIM-3.CAR-CIK cells presented a lower proportion of TIM-3^+^/PD-1^+^ cells at all E:T ratios analyzed compared to non-transduced (NT) cells. To further validate our observations, we also included CD33.CAR-CIK cells in the analysis as a model of antigen-driven activation. Notably, after 7 days of co-culture, CD33.CAR stimulation led to a higher proportion of TIM-3^+^/PD-1^+^ cells than TIM-3.CAR-CIK cells suggesting that the latter may maintain a less exhausted profile upon prolonged antigen exposure (**Figure S2J**). Together, these results demonstrate that TIM-3.CAR-CIK cells selectively recognize and eliminate TIM-3⁺ AML cells, proliferate in an antigen-dependent manner, and produce effector cytokines while maintaining a more favorable exhaustion profile upon chronic antigen encounter.

### TIM-3.CAR-CIK cells recognize and eliminate TIM-3^+^ primary AML blasts and LSCs in vitro

Subsequently, we evaluated whether TIM-3.CAR-CIK cells could target patient-derived AML samples. In long-term co-culture, TIM-3.CAR-CIK cells showed consistent cytotoxic activity against AML blasts compared with control NT cells, demonstrating sustained antileukemic activity (**Figure 2A**). To further investigate antigen specificity, we analyzed the residual percentage of TIM-3^+^ blasts after 7 days of co-culture across different E:T ratios, showing that TIM-3.CAR-CIK cells markedly reduced the frequency of TIM-3^+^ blasts (**Figure 2B**). Because TIM-3 is enriched on LSC compartments, we assessed the impact of TIM-3.CAR-CIK cells on LSC-enriched subsets. TIM-3.CAR-CIK cells efficiently depleted CD34⁺CD38⁻ cells and reduced the frequency of leukemic cells expressing G-protein coupled receptor 56 (GPR56), a novel and stable LSC marker^33^, indicating effective targeting of multiple phenotypically defined LSC populations (**Figures 2C-2D**). Consequently, we observed a target-specific proliferative response of TIM-3.CAR-CIK cells against AML blasts (**Figure 2E**) and increased production of IFN-γ and IL-2 after co-culture with primary AML blasts compared to unstimulated conditions (**Figure 2F**). Together, these data show that TIM-3.CAR-CIK cells selectively eliminate TIM-3⁺ primary AML blasts and LSC-enriched subsets while mounting strong proliferative and cytokine responses.

**Figure 2.**
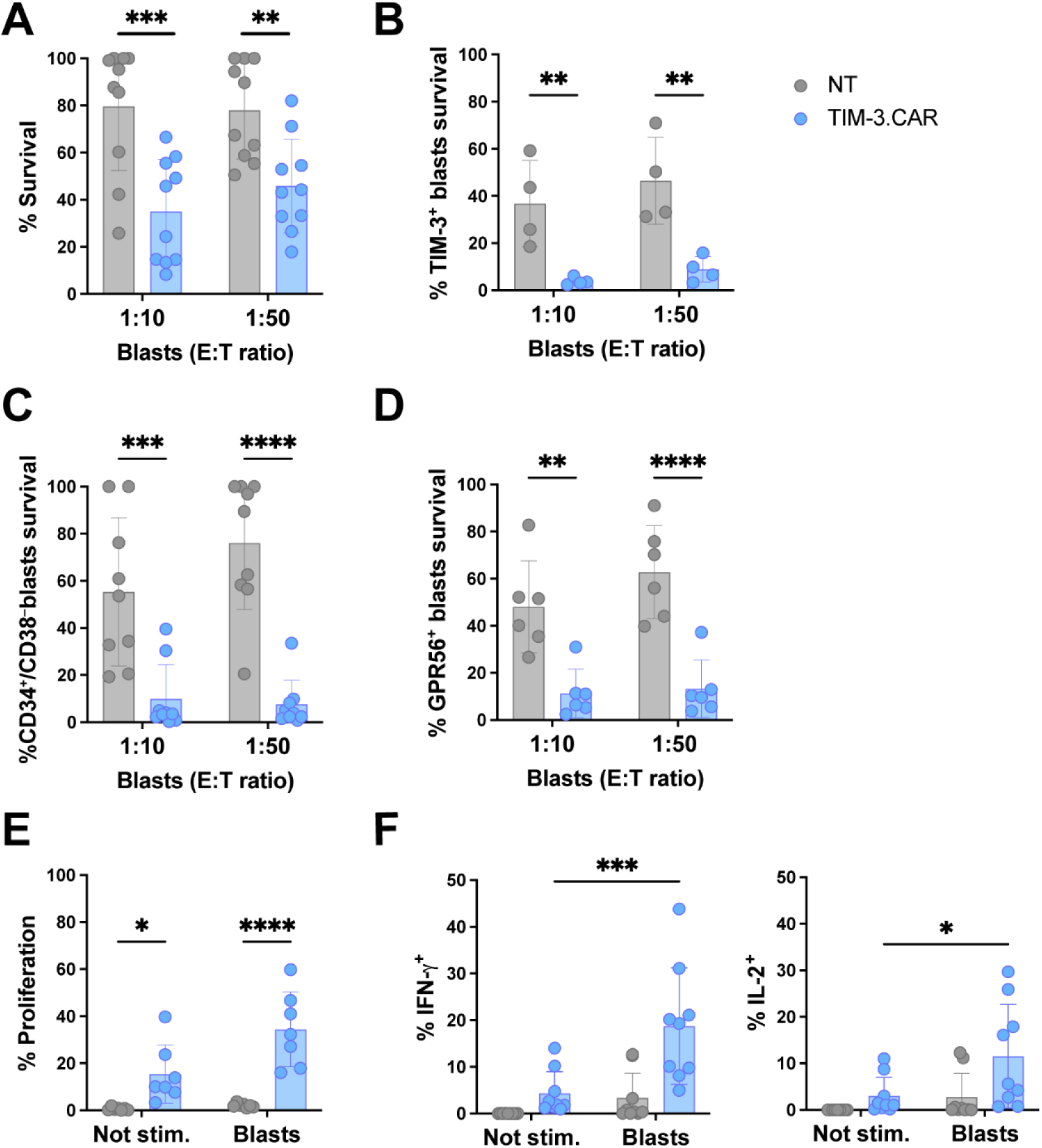
TIM-3.CAR-CIK cells significantly reduce AML blast and LSC survival, while retaining proliferative capacity and effector cytokine production. **(A)** Long-term killing assay of TIM-3.CAR-CIK cells against four primary AML samples compared with NT cells. Blasts survival was assessed by flow cytometry (E:T 1:10 and 1:50, n = 8 donors). See also **Figure S2D**. **(B)** Survival of TIM-3^+^ primary AML blasts (n=3) after 7-day co-culture with TIM-3.CAR-CIK or NT cells (E:T 1:10 and 1:50, n = 4 donors). See also **Figure S2E**. **(C)** Recovery of the LSC-enriched CD34^+^ CD38^-^ population (n = 9) and (**D**) of GPR56^+^ blasts (n = 6) after long-term co-culture. **(E)** Proliferation of TIM-3.CAR-CIK cells assessed by Ki67 staining after 72 hours co-culture with AML blasts (E:T 1:1, n = 7). See also **Figure S2G**. **(F)** Cytokine production (IFN-γ, IL-2) after 5 hours co-culture of TIM-3.CAR-CIK or NT cells with primary AML blasts (E:T 1:3, n = 9). See also **Figure S2H**. Data are presented as individual values and mean ± SD. Statistics were calculated with repeated-measures two-way ANOVA with Bonferroni’s post hoc test. ns, not significant; *p = 0.01, **p < 0.001, ***p = 0.0001 and ****p < 0.0001. See also **Figure S2** for TIM-3.CAR validation in KASUMI-3, AML cell line.

### Minimal off-tumor activity of TIM-3.CAR-CIK cells reflects preferential recognition of AML-enriched, fucosylated TIM-3 glycoforms

TIM-3 is expressed across multiple immune cell types, such as CIK cells, monocytes and NK cells (**Figure 3A**), raising concern for TIM-3.CAR-mediated on-target/off-tumor toxicity. Notably, TIM-3.CAR-CIK cells showed only minimal cytotoxicity against these populations while retaining potent activity against TIM-3⁺ KASUMI-3 cells. Consistently, activated TIM-3⁺ CIK cells were not targeted in an allogeneic setting, and monocytes and NK cells were largely spared (**Figures 3B–3C**). These findings indicate that TIM-3.CAR-CIK cells selectively eliminate leukemic TIM-3⁺ cells while sparing the healthy immune subsets tested, consistent with limited off-tumor reactivity.

**Figure 3.**
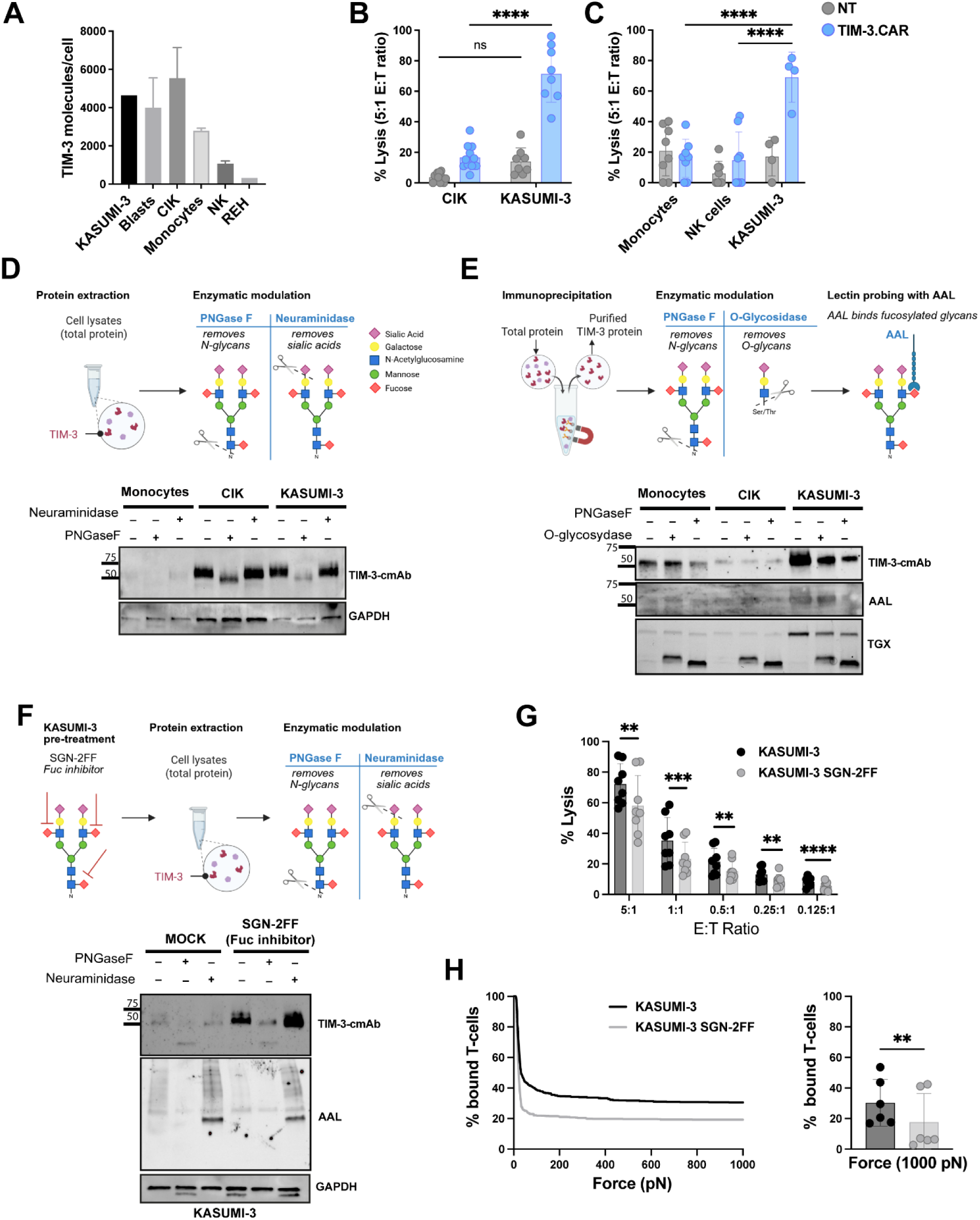
Minimal off-tumor activity of TIM-3.CAR-CIK cells and biochemical evidence for AML-enriched TIM-3 N-glycan features. **(A)** TIM-3 expression on KASUMI-3 cells, primary AML blasts, and healthy immune subsets (CIK cells, monocytes, NK cells) assessed by flow cytometry using QuantiBRITE beads. REH (ALL cell line) served as negative control. **(B)** Short-term killing assay of TIM-3.CAR-CIK cells against CIK (n = 11) or KASUMI-3 (n = 8) cells compared with NT cells. Target cell lysis was evaluated by flow cytometry (E:T 5:1). **(C)** Short-term killing assay of TIM-3.CAR-CIK cells against monocytes (n = 8) or NK cells (n = 8) compared with NT (E:T 5:1). KASUMI-3 (n = 4) were included as positive control. **(D)** Immunoblot analysis of TIM-3 in lysates from monocytes, CIK cells, and KASUMI-3 cells following enzymatic treatment with PNGase F or broad neuraminidase, probed with a commercial anti–TIM-3 antibody (TIM-3-cmAb). GAPDH, loading control. Glycan symbols follow SNFG. **(E)** TIM-3 immunoprecipitates from monocytes, CIK cells, and KASUMI-3 cells treated with PNGase F or *O-*glycosidase and analyzed by immunoblot with TIM-3-cmAb and lectin far-western with Aleuria aurantia lectin (AAL; fucosylated epitopes). TGX stain-free total protein signal is shown as a loading/normalization control. **(F)** KASUMI-3 cells treated with vehicle (mock) or the fucosylation inhibitor 2F-peracetyl-fucose (SGN-2FF), followed by PNGase F or neuraminidase treatment and immunoblot/lectin probing with TIM-3-cmAb and AAL. See also **Figure S3A**. **(G)** Short-term killing assay of TIM-3.CAR-CIK cells against untreated or SGN-2FF-treated KASUMI-3 cells at various E:T ratios (5:1, 1:1, 0.5:1, 0.25:1 and 0.125:1, n = 8). **(H)** Affinity kinetics (left) and binding avidity at 1000 pN force (right) of TIM-3.CAR-CIK cells to untreated or defucosylated KASUMI-3 by LUMICKS analysis (n = 6). Immunoblot experiments (D-F) were repeated in three independent biological replicates with similar results. Data are presented as individual values and mean ± SD. Statistical significance was determined with repeated-measures two-way ANOVA with Bonferroni’s post hoc test (B, C) or using paired t test (G, H). ns, not significant; *p = 0.01, **p < 0.001, ***p = 0.0001 and ****p < 0.0001. Illustrations were created with Biorender.com. See also **Figure S3** for loading-matched TIM-3 immunoprecipitation controls.

We therefore investigated alternative mechanisms underlying the observed selective tumor targeting. Aberrant tumor-specific glycosylation is a well-established hallmark of malignancy and can generate tumor-restricted epitopes exploitable for immunotherapy^34,35^. In particular, terminal fucosylation and hyper-sialylation are pervasive cancer-associated glycan alterations that can modulate recognition of cell-surface glycoproteins^36,37^. Given the dense glycosylation of the TIM-3 mucin domain^38,39^, we explored whether TIM-3 in AML carries distinct glycoforms that could contribute to preferential TIM-3.CAR recognition relative to healthy immune cells.

We first assessed TIM-3 protein abundance and glycosylation status in monocytes, CIK cells, and KASUMI-3 cells by immunoblot. Using a commercial anti-TIM-3 antibody (TIM-3-cmAb), we confirmed high TIM-3 expression in CIK and KASUMI-3 cells (**Figure 3D**), consistent with prior surface-expression patterns (**Figure 3A**). Treatment with the *N*-glycan cleaving enzyme PNGase F, caused a clear downward mobility shift of the TIM-3 band across all tested cell types, indicating that TIM-3 carries *N-*linked glycans. This is consistent with the single UniProt-annotated *N-*glycosylation site (Asn172; UniProt Q8TDQ0) in human TIM-3 (HAVCR2). Notably, PNGase F treatment markedly reduced TIM-3-cmAb signal intensity, suggesting glycosylation-sensitive epitope detection under these blotting conditions. In contrast, broad neuraminidase treatment produced no obvious change in TIM-3 detection in this lysate immunoblot setup, suggesting that terminal sialic acids are not rate-limiting for TIM-3-cmAb binding (**Figure 3D**). Together, these data establish that TIM-3 is *N-*glycosylated in monocytes, CIK, and KASUMI-3 cells and highlight an *N-*glycan–dependent component to TIM-3 detection by TIM-3-cmAb.

To further dissect TIM-3 glycosylation across cell types, we immunoprecipitated TIM-3 and assessed immunoreactivity following PNGase F or *O-*glycosidase treatment, alongside lectin far-western probing with Aleuria aurantia lectin (AAL) to detect fucosylated glycans. In KASUMI-3 cells, TIM-3-cmAb detection was strongly reduced by PNGase F and only modestly affected by *O-*glycosidase (**Figure 3E**), indicating predominant dependence on *N-*linked glycans. Consistently, AAL binding was largely preserved after *O-*glycosidase digestion but markedly decreased upon PNGase F treatment in KASUMI-3 cells, supporting that AAL-reactive fucosylated epitopes are mainly carried on TIM-3 *N-*glycans in AML cells. By contrast, in monocytes and CIK cells, TIM-3-cmAb detection was also reduced by PNGase F, but to a lesser extent than in KASUMI-3 cells, and AAL signal changes were similarly more modest under the same conditions. These findings are consistent with a lower abundance of AAL-reactive fucosylated *N-*glycans on TIM-3 in healthy immune subsets. Consistent with glycan-dependent masking of the TIM-3-cmAb epitope, acute defucosylation with the fucosyltransferase inhibitor 2F-peracetyl-fucose (SGN-2FF) increased TIM-3-cmAb detection in KASUMI-3 cells, accompanied by the expected reduction in AAL reactivity on the same blot. Neuraminidase-mediated desialylation further enhanced TIM-3-cmAb detection, suggesting that terminal sialylation can further limit epitope accessibility and reduce antibody recognition (**Figure 3F**). Notably, under matched loading conditions (**Figure S3A**), TIM-3-cmAb reactivity remained the lowest for KASUMI-3-derived TIM-3, despite robust expression, confirming that differential detection reflects glycan composition rather than antigen abundance. Together, these data indicate that TIM-3 on KASUMI-3 carries distinct *N-*glycan features, consistent with increased fucosylation and sialylation relative to healthy immune cells.

We next tested whether the AML-enriched fucosylation signature functionally contributes to productive TIM-3.CAR engagement. To address this, we inhibited cellular fucosylation in KASUMI-3 cells using SGN-2FF^35^ and assessed CAR-mediated activity. SGN-2FF pre-treatment significantly impaired TIM-3.CAR-CIK recognition and cytotoxicity across multiple effector-to-target ratios (**Figure 3G**), indicating that fucose-containing glycan features are required for efficient CAR-mediated killing. Consistent with this functional impairment, single-cell force spectroscopy (z-Movi; LUMICKS) revealed reduced interaction kinetics (**Figure 3H, left**) and markedly decreased binding avidity at 1000 pN (**Figure 3H, right**) between TIM-3.CAR-CIK cells and defucosylated targets compared with untreated KASUMI-3 cells. Importantly, these effects were observed at the level of CAR-mediated function and interaction strength, providing a direct readout of surface engagement beyond biochemical analyses. Together, these data provide direct functional evidence that fucosylated glycan features on AML TIM-3 enhance CAR–target interaction strength and support productive CAR engagement leading to cytotoxicity.

### Mechanistic basis of glycoform-selective TIM-3.CAR engagement: differential N-glycan remodeling and terminal capping in AML versus healthy immune cells

To test whether the TIM-3.CAR scFv preferentially engages AML-associated TIM-3 glycoforms, we generated a recombinant scFv-derived monoclonal antibody (TIM-3scFv-mAb) as a surrogate for CAR epitope probing. In monocytes, CIK cells, and KASUMI-3 lysates, TIM-3scFv-mAb detected TIM-3 across a broad apparent molecular-weight range. Notably, in untreated KASUMI-3 samples, TIM-3scFv-mAb preferentially recognized the upper high-molecular-weight (HMW) fraction, whereas in monocytes and CIK lysates, it predominantly detected lower-mass TIM-3 species, consistent with distinct TIM-3 glycoform distributions. Upon PNGase F digestion and neuraminidase treatment, TIM-3 species shifted to lower apparent molecular weights as *N-*glycans were removed, yet TIM-3scFv-mAb binding was preserved and tracked the mobility shifts (**Figure 4A**), consistent with an epitope that remains accessible to the TIM-3.CAR scFv after enzymatic glycan trimming in this immunoblot context.

**Figure 4.**
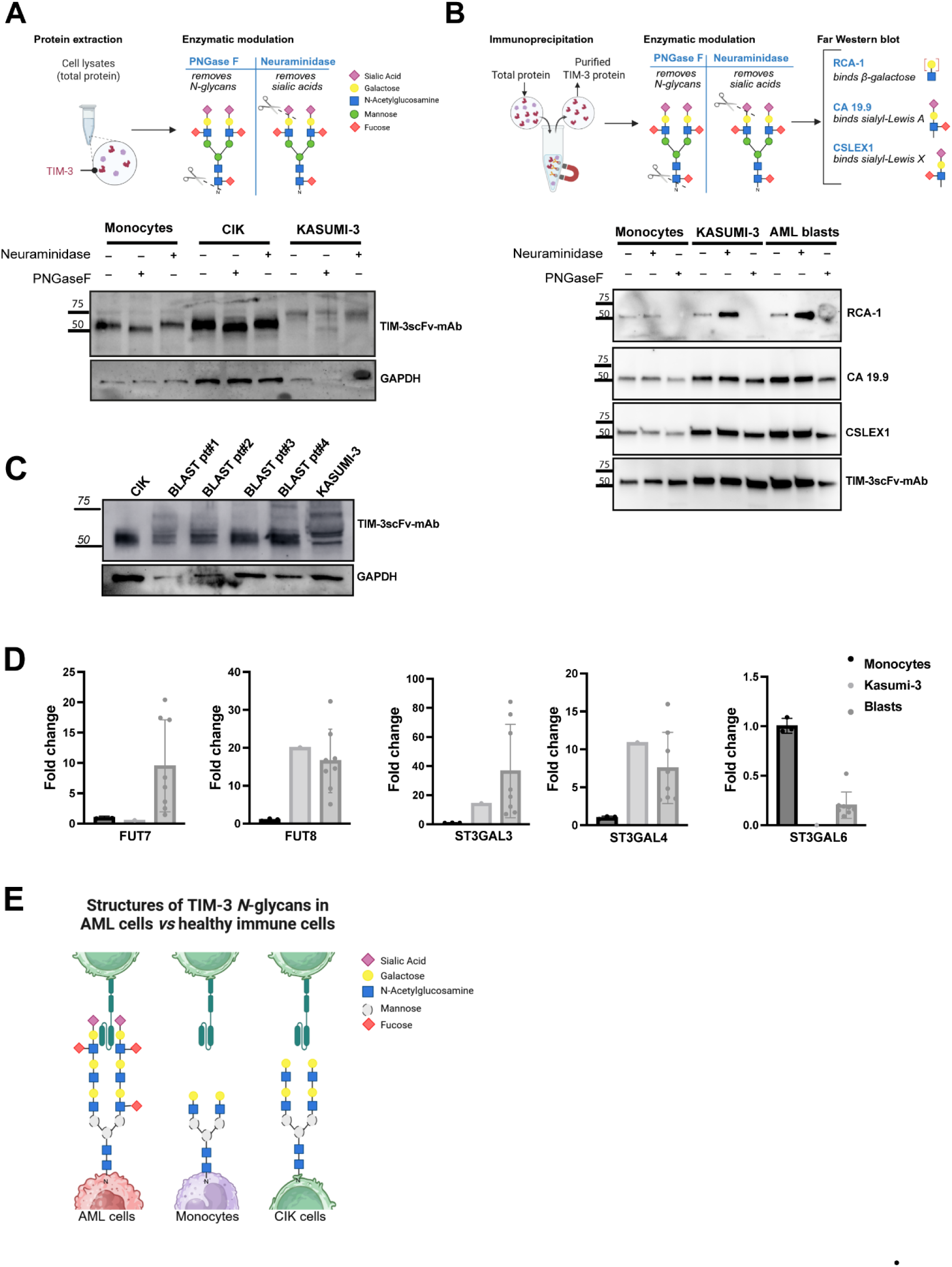
TIM-3.CAR scFv probing identifies AML-associated TIM-3 glycoforms and terminal sialylation/Lewis-type features. **(A)** Immunoblot profiling of TIM-3 glycoforms in monocytes, CIK cells, and KASUMI-3 lysates using a recombinant scFv-derived monoclonal antibody (TIM-3scFv-mAb) following enzymatic treatment with PNGase F or broad neuraminidase. GAPDH, loading control. **(B)** TIM-3 immunoprecipitates from healthy monocytes, KASUMI-3 cells, and primary AML blasts treated with neuraminidase and/or PNGase F and analyzed by lectin and antibody probing: Ricinus communis agglutinin I (RCA-I; terminal β-galactose/LacNAc motifs), CA19-9 (sialyl-Lewis A), CSLEX1 (sialyl-Lewis X), and TIM-3scFv-mAb. See also **Figure S3B**. **(C)** High-resolution immunoblot of TIM-3 species detected by TIM-3scFv-mAb in CIK cells, primary AML blasts, and KASUMI-3 cells. GAPDH, loading control. See also **Figure S3C**. **(D)** RT-qPCR expression profiling of glycosyltransferases (FUT7, FUT8, ST3GAL3, ST3GAL4, ST3GAL6) in monocytes, KASUMI-3 cells, and primary AML blasts. Data are plotted as fold-change relative to monocytes and normalized to 18S RNA; individual points denote biological samples where applicable. **(E)** Schematic model summarizing a glycoform-biased recognition framework in which AML-associated remodeling of TIM-3 *N*-glycans contributes to preferential TIM-3.CAR recognition of AML-enriched TIM-3 glycoforms. Representative *N*-glycan structures are proposed for TIM-3 in AML blasts, monocytes and CIK cells based on enzymatic perturbation and lectin/antibody probing. Sugar moieties drawn with dashed outlines indicate features not directly resolved/assigned. Glycan symbols follow SNFG. Immunoblot and lectin/antibody blot experiments (A-C) were repeated in three independent biological replicates with similar results. Illustrations were created with Biorender.com. See also **Figure S3** for additional lectin/antibody probing of TIM-3 glycoforms and terminal galactose exposure.

We next asked whether AML-associated alterations in TIM-3 hyper-glycosylation include specific terminal glycan motifs such as sialyl–Lewis–type determinants, which are frequently enriched in AML and associated with aberrant fucosylation and sialylation pathways^40^. Because TIM-3 contains a mucin-rich ectodomain with high glycan density, efficient enzymatic removal of *O-*linked glycans can be technically challenging^39^. Nevertheless, probing TIM-3 immunoprecipitated with CSLEX1 (sialyl-Lewis X) showed sensitivity to PNGase F (**Figure S3B**), suggesting that a fraction of these determinants is carried on *N-*linked glycans. Extending this analysis to TIM-3 immunoprecipitated from healthy monocytes, KASUMI-3 cells, and primary AML blasts, we probed TIM-3 glycan termini by far-western blot using *Ricinus communis* agglutinin 1 (RCA-1), a tetrameric lectin that preferentially binds terminal β-galactose, particularly within LacNAc motifs (Galβ1-4GlcNAc)^41,42^. Neuraminidase treatment markedly increased RCA-1 binding to TIM-3 in KASUMI-3 cells and AML blasts relative to monocytes (**Figure 4B**). Because neuraminidase selectively removes terminal sialic acids, this increase is consistent with desialylation exposing underlying RCA-1–reactive β-galactose/LacNAc termini that were previously capped by sialic acids. Across cell types, RCA-1 reactivity increased from monocytes to CIK cells to KASUMI-3, was further enhanced by neuraminidase, and was abolished by PNGase F (**Figure S3C**), supporting cell-type–dependent differences in the exposure of terminal β-galactose/LacNAc motifs on TIM-3 N-glycans. Together, the neuraminidase-unmasked RCA-1 signal is consistent with an enriched exposure of β-galactose/LacNAc termini on TIM-3 *N-*glycans in KASUMI-3 cells and AML blasts. Importantly, PNGase F abrogated RCA-1 reactivity, indicating that these neuraminidase-unmasked termini reside predominantly on *N-*linked glycans (**Figure 4B**). In parallel, CSLEX1 and CA19-9 (recognizing sialyl-Lewis X and sialyl-Lewis A, respectively) signals decreased upon desialylation and were reduced by PNGase F, supporting the presence of sialyl-Lewis–type determinants on a subset of TIM-3 *N*-glycans in KASUMI-3 cells and AML blasts (**Figure 4B**). Consistent with these glycoform features, AML blasts displayed prominent high-molecular-weight TIM-3 species resembling KASUMI-3 cells (**Figure 4C**).

Moreover, RT–qPCR profiling across monocytes, KASUMI-3 cells, and primary AML blasts supported an AML-skewed glycosyltransferase program (**Figure 4D**). In KASUMI-3, FUT8 was strongly upregulated (∼20× over monocytes), together with ST3GAL3 (∼14×) and ST3GAL4 (∼11×), consistent with increased capacity for core fucosylation and α2,3-sialylation. In primary AML blasts, we observed substantial inter-sample variability but an overall elevation of ST3GAL3 (mean ∼36×; range ∼7–84×), along with increased FUT8 (mean ∼17×; range ∼5–32×), FUT7 (mean ∼9.5×; range ∼1.5–20×), and ST3GAL4 (mean ∼7.5×; range ∼3.3–16×). By contrast, ST3GAL6 remained low in both KASUMI-3 and AML blasts (≤∼0.5×), suggesting that α2,3-sialylation in this context is more consistent with ST3GAL3/4 than ST3GAL6 at the transcript level. This expression signature is coherent with the biochemical and glycan-probing data, including enhanced AAL reactivity, neuraminidase-unmasked RCA-I binding, and the presence of sialyl-Lewis–type determinants on TIM-3 immunoprecipitated from AML samples (**Figures 3E–3F and 4B–4C**).

Together, these data support a model in which AML cells present TIM-3 glycoforms enriched in terminal fucosylation and sialylated/Lewis-type features, consistent with an AML-biased expression of glycosylation enzymes. While TIM-3scFv-mAb retains detection of enzymatically remodeled TIM-3 species in immunoblots (**Figure 4A**), functional perturbation of fucosylation reduces CAR–target binding avidity and cytotoxicity (**Figures 3G–3H**), suggesting that terminal *N-*glycan features enhance productive and stable TIM-3.CAR engagement on AML targets. This glycoform-biased recognition model is summarized in **Figures 4E**, which provides a mechanistic framework for potent anti-AML activity with limited off-tumor reactivity despite TIM-3 expression on healthy immune subsets.

### TIM-3.CAR-CIK cells display efficient control of AML burden in vivo

To evaluate the *in vivo* antileukemic activity of TIM-3.CAR-CIK cells, we established a leukemia xenograft model. Sublethally irradiated NSG mice were inoculated with 10^7^ KASUMI-3 cells (day 0) and subsequently treated with three weekly administrations of TIM-3.CAR-CIKs (10^7^ cells/ mouse on days 14, 21, 28) or left untreated (CTR). Leukemic burden was assessed on day 35 by quantifying hCD45⁺CD33⁺ and hCD45⁺TIM-3⁺ cells in bone marrow (BM), spleen, and peripheral blood (PB). Because KASUMI-3 cells are not uniformly TIM-3⁺, residual leukemic cells were primarily analyzed by CD33⁺ expression (100% positive), while TIM-3 analysis was performed to confirm CAR specificity (**Figure 5A**). Mice treated with TIM-3.CAR-CIK cells exhibited a significant, though incomplete, reduction of hCD33⁺ cells in the BM (CTR mean 81.5% vs. 53.2% in treated mice, p = 0.0013) as well as decreased frequencies of hTIM-3⁺ cells (CTR mean 61.5% vs. 40.5% in treated mice, p = 0.0167) (**Figure 5B–C**). A comparable reduction in leukemic burden was observed in the spleen (**Figure 5D**), and circulating hCD33⁺ and hTIM-3⁺ cells in PB were markedly lower in treated animals compared with controls (**Figure 5E**).

**Figure 5.**
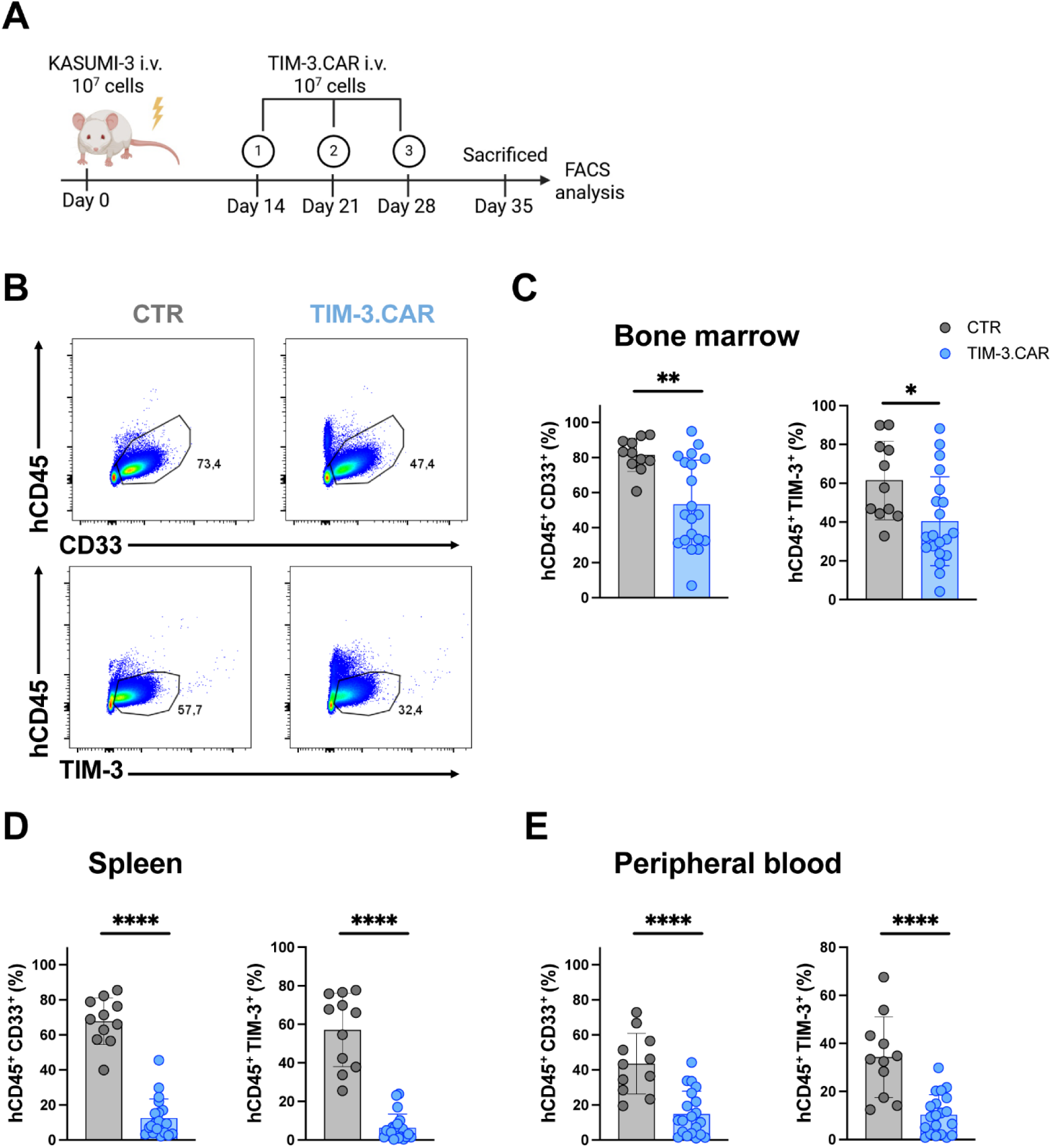
TIM-3.CAR-CIK treatment abates TIM-3^+^ KASUMI-3 leukemia burden in NSG mice. **(A)** Schematic of the xenograft KASUMI-3 AML model. **(B)** Representative flow cytometry plots of hCD45^+^CD33^+^ (up) and of hCD45^+^TIM-3^+^ cells (down) in the BM of CTR or TIM-3.CAR treated mice at sacrifice. (**C-E**) Frequencies of hCD33^+^ and hTIM-3^+^ cells in the (C) BM, (D), spleen and (E) peripheral blood (PB) at sacrifice. Illustrations were created with Biorender.com. Results represent three independent experiments using TIM-3.CAR-CIK cells generated from 3 different donors. Data are presented as individual values and mean ± SD. Statistical significance was determined by unpaired t test. *p = 0.01, **p < 0.001 and ****p < 0.0001.

Collectively, these results demonstrate that TIM-3.CAR-CIK cells exert significant antileukemic activity *in vivo*, reducing AML burden despite heterogeneous TIM-3 expression on target cells.

### CIK cells can be efficiently engineered for IF-BETTER dual targeting of TIM-3 and CD33, resulting in effective in vitro elimination of AML blasts and LSCs

Since TIM-3.CAR-CIK cells significantly reduced, but did not fully eradicate, BM leukemic burden *in vivo*, likely due to heterogenous TIM-3 expression among AML cells, we explored a dual-targeting strategy. We therefore designed next-generation CAR constructs, pairing TIM-3 with CD33 and exploiting the IF-BETTER gating approach (**Figure 6A**). To validate target selection, we analyzed the co-distribution of TIM-3 and CD33 in a cohort of newly diagnosed pediatric and adult AML patients with diverse genetic and molecular backgrounds (n=44), comparing bulk AML blasts with LSC-enriched populations (CD34^+^CD38^-^). Antigens densities were overall heterogeneous, with increased CD33^high^ density on bulk and TIM-3^high^ density on both blasts and LSCs, further confirming TIM-3 as a specific LSC marker^14,16^ (**Figure 6B**) .Two Dual CAR constructs were generated, each composed of a second-generation CAR paired with a CCR, recognizing CD33 and TIM-3, in the two configurations: CD33.CAR/TIM-3.CCR and TIM-3.CAR/CD33.CCR. The anti-CD33 scFv was derived from high-affinity, humanized rat anti-human CD33 mAb, while the anti-TIM-3 scFv was the one previously used in the single-target CAR (**Figure 6C**). Dual CAR-CIK cells were successfully generated following the above-mentioned protocol, showing at the end of the culture an overall expression of 60% (**Figure S4A**) and a remarkable co-expression of CAR and CCR molecules (n = 18 different donors) (**Figure S4B**). The transduction process did not affect the typical CIK phenotype, the distribution of CD4 and CD8 subpopulations or the memory status of CAR-CIK cells, confirming the suitability of SB-based delivery system (**Figure S4C-E**). Moreover, we analyzed the fold increase of Dual CAR-CIK cells at early time points during culture period to validate the absence of fratricide effect, as previously assessed for TIM-3.CAR-CIK cells, and to investigate any potential toxicity associated with a longer CAR sequence integration. Notably, both CD33.CAR/TIM-3.CCR and TIM-3.CAR/CD33.CCR configurations expanded comparably to single CD33.CAR and TIM-3.CAR-CIK cells, confirming that no fratricide nor DNA toxicity occurred (**Figure S4F**).

**Figure 6.**
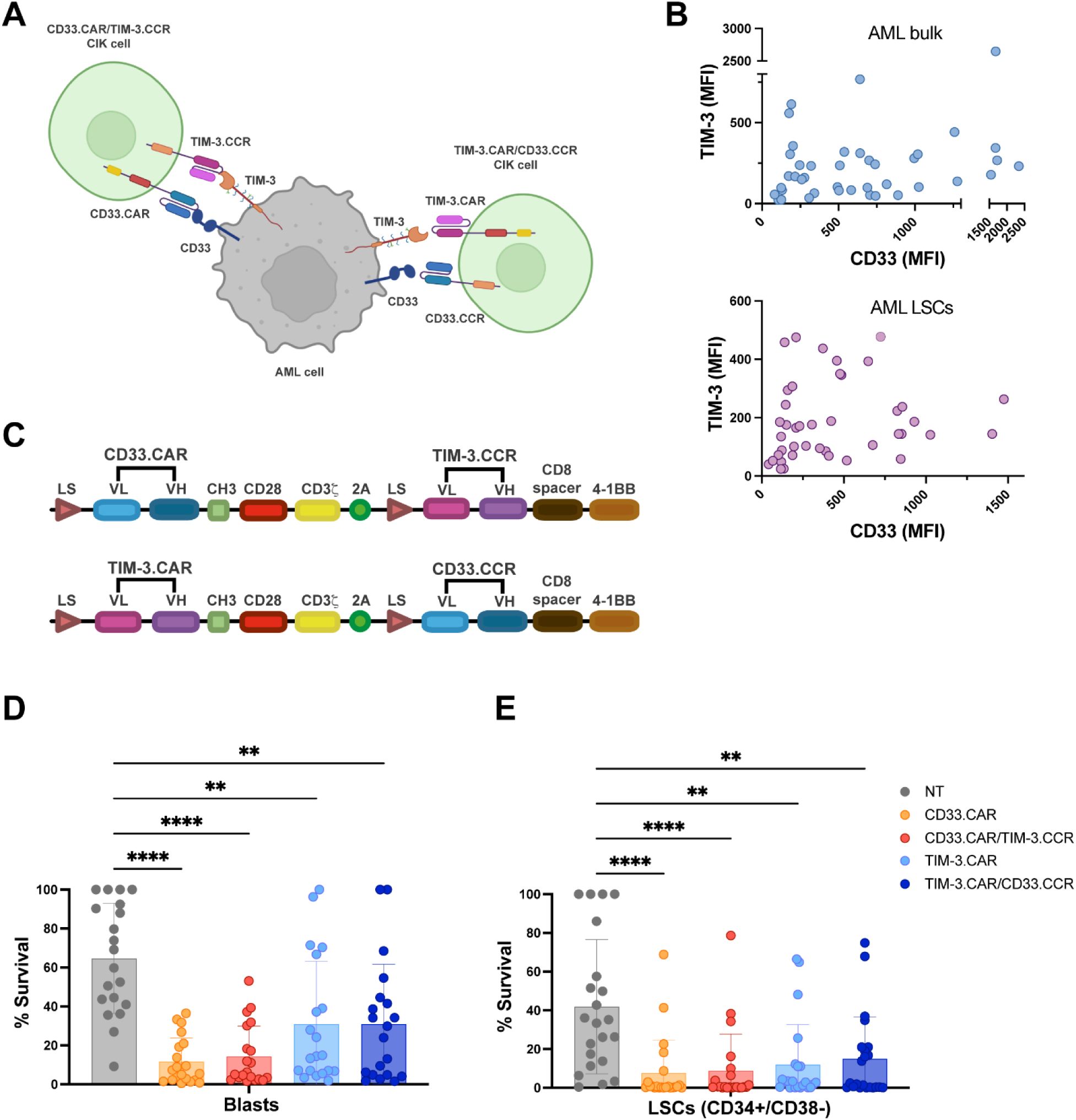
IF-BETTER dual targeting CD33.CAR/TIM-3.CCR and TIM-3.CAR/CD33.CCR constructs are effectively expressed and functionally active in CIK cells. **(A)** Schematic of IF-BETTER gate strategy showing dual antigen recognition of CD33^+^/TIM-3^+^ target cell by CD33.CAR/TIM-3.CCR and TIM3.CAR/CD33.CCR-CIK cells. **(B)** Co-distribution of CD33 and TIM-3 expression (MFI) on bulk AML (top) and LSC-enriched CD34^+^CD38^-^population (bottom). Each dot represents a distinct patient (n=44 patients). **(C)** Schematics of next-generation Dual CD33.CAR/TIM-3.CCR and TIM-3.CAR/CD33.CCR constructs. CAR molecules are second-generation, carrying CD28 co-stimulatory domain, while CCR molecules present 4-1BB as co-stimulus. Both constructs were cloned into a pT4-transposon vector. See also **Figure S4A, B**. **(D)** Long-term killing assay (E:T 1:10) of all CAR-CIK cells against primary AML blasts (n=8 blasts) compared to NT cells. Blasts survival was determined by flow cytometry (n=13 donors). **(E)** Recovery of LSC-enriched CD34^+^CD38^-^ population (n=8 patient samples) after 7 days co-culture with all CAR-CIK cells (E:T 1:10), compared to NT cells (n=13). Data are presented as individual values and the mean ± SD. Statistical significance was determined by one-way ANOVA test. ** p<0.01, **** p<0.0001. Illustrations were created with Biorender.com. See also **Figure S4** for expression and phenotypic characterization of Dual CAR constructs, **Figure S5** for Dual CAR-CIK cell activity against AML cell lines and **Figure S6** for Dual CAR-CIK cell off-tumor toxicity against healthy immune and hematopoietic cells.

To assess Dual CAR-CIK cells effector functions, we selected KG-1 cells (CD33^high^TIM-3^low^) and engineered a TIM-3^high^ KG-1 variant, establishing a CD33^high^TIM-3^high^ AML model to test both Dual CARs across different antigen densities. REH cell line (CD33^-^TIM-3^-^) was used as negative control (**Figure S5A**). When tested in a short-term cytotoxic assay, CD33.CAR/TIM-3.CCR-CIK cells mediated significantly higher lysis of KG-1 cells (mean: 66.2%) compared with TIM-3.CAR (mean: 38%, n = 10, p = 0.0013) and TIM-3.CAR/CD33.CCR-CIK cells (mean: 36.8%, n = 10, p = 0.0008) consistent with the low TIM-3 expression on the parental line. Both Dual CAR constructs displayed efficient cytotoxic activity against KG-1 TIM-3^+^ cells, indicating that CCR engagement boosts killing of double-antigen positive target cells. No relevant lysis was observed against REH, confirming target specificity (**Figure S5B**). Furthermore, both CD33.CAR-/TIM-3.CCR and TIM-3.CAR/CD33.CCR-CIK cells secreted significantly higher levels of IFN-γ and IL-2 in response to KG-1 compared with TIM-3.CAR-CIK cells (n = 10) further supporting the role of CCR engagement when the CAR target antigen is expressed at low levels. Notably, TIM-3.CAR/CD33.CAR-CIK cells showed the highest IL-2 production in response to KG-1 TIM-3^+^ cells (mean: 31,7%, n = 3) compared with parental KG-1 (mean: 20.4%, n = 10) and REH (mean: 1.35%, n = 7), suggesting strong costimulatory signaling via CD33 (**Figure S5C**). Overall, these results suggest that IF-BETTER Dual CAR-CIK cells effectively kill AML cells with different CD33 and TIM-3 antigen densities, offering a suitable alternative to the previous system based on single-target TIM-3.CAR-CIK cells.

Afterwards, we tested the efficacy of Dual CAR-CIK cells *in vitro* against patient-derived AML blasts, exhibiting variable CD33 and TIM-3 expression (n=8 patients). In long-term cytotoxicity assays (E:T ratio = 1:10), both CD33.CAR/TIM-3.CCR and TIM-3.CAR/CD33.CCR-CIK cells effectively eradicated leukemic blasts, with a number of retrieved blasts comparable to their single CAR-CIK cells counterpart (**Figure 6D**). We also analyzed the percentage of CD34^+^CD38^-^ cells retrieved after the long-term co-culture, observing that both CD33.CAR/TIM-3.CCR and TIM-3.CAR/CD33.CCR-CIK cells were able to kill the CD34^+^CD38^-^subset, thus confirming their capability to effectively eliminate LSC-enriched populations (**Figure 6E**).

Together, these findings show that IF-BETTER Dual CAR-CIK cells can be efficiently engineered and provide a potent, flexible approach to eliminate AML blasts and LSCs across heterogeneous antigen-expression states.

### CD33.CAR/TIM-3.CCR and TIM-3.CAR/CD33.CCR-CIK cells spare healthy differentiated CD33^+^/TIM-3^+^ cells and HSCs in vitro

Since we observed that TIM-3.CAR-CIK cells display minimal toxicity against healthy immune TIM-3^+^ cells (**Figure 3B-C**) and the CD33.CAR-CIK cells on-target/off-tumor toxicity is extensively documented^43,44^, we wanted to confirm the safety profile also for CD33.CAR/TIM-3.CCR and TIM-3.CAR/CD33.CCR-CIK cells. We first evaluated the surface expression of CD33 and TIM-3 on healthy donor monocytes and NK cells (**Figure S6A**). Next, to determine whether Dual CAR-CIK cells mediate off-target toxicity, we performed short-term *in vitro* killing assays. As expected, both Dual CAR-CIK cells exhibited minimal killing activity against monocytes, NK cells and activated T cells, and a much superior lysis against KG-1 (**Figure S6B**). Moreover, we assessed the potential hematotoxicity of Dual CAR-CIK cells by co-culturing them with HSCs and subsequently seeding the remaining cells in a Colony Forming Unit (CFU) assay. Notably, HSC colony-forming ability was preserved, yielding a number of colonies comparable to NT cells (**Figure S6C**).

### Dual CAR-CIK cells targeting CD33 and TIM-3 effectively control AML burden in vivo

To investigate the therapeutic potential of Dual CAR-CIK cells *in vivo*, we established both low- and high-burden leukemic xenograft mouse models using a schedule which consists on the infusion of KG-1 TIM-3^+^ (CD33^high^TIM-3^high^) AML cell line (5×10^6^ cells/mouse) and subsequent treatment, at two different time points, with TIM-3.CAR, CD33.CAR/TIM-3.CCR, or TIM-3.CAR/CD33.CCR-CIK cells (10^7^ cells/mouse).

In the low-burden setting mice received CAR-CIK infusions on day 5 and 12 after leukemia inoculation (**Figure 7A**). In the BM, both Dual CAR configurations markedly reduced hCD45⁺CD33⁺ leukemic cells compared with TIM-3.CAR-CIK cells (**Figure 7B-C**). TIM-3 expression on residual hCD45^+^ CD33^+^ leukemic cells in the BM revealed that TIM-3^+^ cells were effectively targeted in all treatment groups compared to controls. (**Figure 7D**). Leukemic burden in spleen and PB was similarly decreased, with Dual CAR-CIK cells showing the most profound clearance (**Figure 7E**).

**Figure 7.**
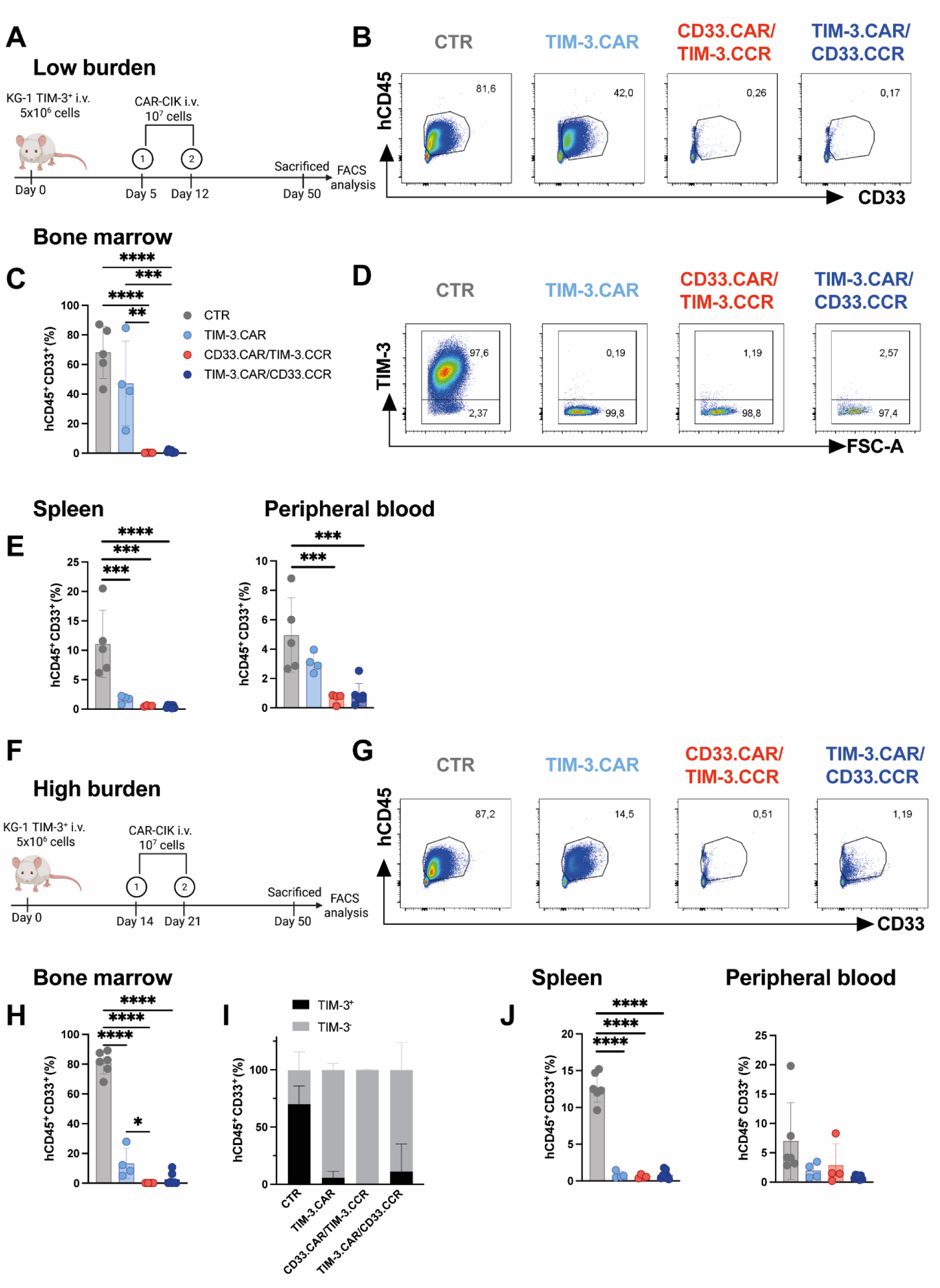
CD33.CAR/TIM-3.CCR and TIM-3.CAR/CD33.CCR-CIK cells display improved clearance of KG-1 TIM-3^+^ leukemia burden in NSG mice. (**A, F**) Schematic of KG-1 TIM-3^+^ low (A) and high (F) burden *in vivo* experimental design. (**B, G**) Representative FACS plots of hCD45^+^CD33^+^ cells in the BM of CTR and CAR-treated mice in low-(B) and high-burden (G) settings. (**C, H**) Summary of BM hCD33^+^ cell percentages for all groups. (**D, I**) Analysis of TIM-3 expression on residual hCD33^+^ cells in BM of CTR and treated mice. (**E, J**) Leukemic burden in spleen (left) and PB (right) evaluated by flow cytometry, in low- (E) and (J) high-burden models. Illustrations were created with Biorender.com. Results represent two independent experiments using CAR-CIK cells generated from 2 donors. Data are presented as individual values and the mean ± SD. Statistical significance was determined by one-way ANOVA. **p < 0.001, ***p = 0.0001 and ****p < 0.0001.

We next tested Dual CAR-CIK cells in a high-burden model, administering CAR-CIK treatment at day 14 and day 21 after KG-1 TIM-3^+^ injection (**Figure 7F**). TIM-3.CAR-CIK cells significantly reduced BM leukemia but did not achieve full eradication. In contrast, both Dual CAR groups nearly eliminated hCD45⁺CD33⁺ cells in the BM (**Figure 7G-H**), with CD33.CAR/TIM-3.CCR showing significantly greater activity than the single TIM-3.CAR. TIM-3 expression on the few remaining leukemic cells revealed efficient depletion of TIM-3⁺ disease in all CAR-treated mice (**Figure 7I**). Spleen and PB analyses consistently demonstrated robust leukemia control across all compartments (**Figure 7J**).

Together, these findings demonstrate that IF-BETTER Dual CAR-CIK cells provide superior control of AML compared with single-target TIM-3.CAR-CIK cells, achieving near-complete elimination of leukemia in both low- and high-disease-burden settings.

## DISCUSSION

In this work, we address key challenges constraining the therapeutic potential of CAR-T cells in AML, including antigenic heterogeneity, incomplete leukemia eradication, and risk of on-target/off-tumor toxicity. The primary hurdle is the identification of functionally relevant, tumor-specific target antigens, as most AML targets are frequently shared with healthy hematopoietic and immune cells. TIM-3 represents a biologically and therapeutically relevant AML antigen, selectively enriched on LSCs^14,15^ and implicated in leukemic self-renewal^16,21,45^ and immune suppression^22^. Consistently, multiple anti-TIM-3 monoclonal antibodies have advanced into early-phase clinical trials, supporting its translational relevance (NCT02817633, NCT02608268, and NCT03099109). Here, we demonstrate that AML-specific *N*-glycosylation generates a tumor-restricted TIM-3 glycoform that increases the binding avidity of a TIM-3.CAR scFv to its protein-proximal epitope. Incorporation of this glycoform-sensitive TIM-3 scFv into an IF-BETTER dual-antigen platform (CD33.CAR/TIM-3.CCR and TIM-3.CAR/CD33.CCR) further enhances functional selectivity, mitigates potential on-target/off-tumor toxicity, and broadens coverage across AML heterogeneity.

Although TIM-3 has been validated as a promising AML target antigen and explored in multiple CAR designs^46–49^, none have addressed the molecular basis enabling selective recognition of AML cells over healthy TIM-3^+^ immune cells, including monocytes and activated NK and T cells^17^. Notably, despite the broad expression of TIM-3, significant on-target/off-tumor toxicity has not been observed, either in our functional assays or in previous studies, suggesting the existence of AML-specific features contributing to tumor-restricted recognition.

Tumor-specific glycosylation is increasingly recognized as a source of neo-epitopes or exposed structures that can be selectively recognized by CAR-T cells^40,50–52^. Pioneering work by Posey et al, showed that CAR-T cells targeting a common tumor-aberrant *O*-glycosylation on MUC1 specifically recognize multiple solid tumors and mediate *in vivo* anti-tumor efficacy, establishing a paradigm for leveraging aberrant glycosylation as a target for immunotherapeutic exploitation^35^. However, such strategies have rarely been explored in hematological malignancies^53^, despite mounting evidence that AML undergoes extensive remodeling of sialylation and fucosylation^54,55^.

The TIM-3 mucin domain is physiologically highly enriched in glycosylation sites which are poorly characterized in AML^39,56^. Glycan composition and sialylation can impact on epitope accessibility, scFv binding and CAR activation thresholds^35,57,58^, while also shaping receptor signaling and interactions with immunomodulatory galectins^40,59,60^. Among these, TIM-3/Gal-9 axis is upregulated in advanced myeloid disease and associated with poor clinical outcomes^61^. These observations suggest TIM-3 is not just a selective AML-surface marker but an active player of an immunosuppressive signaling network. Indeed, TIM-3 is a well-known checkpoint molecule^17,62^ and our TIM-3.CAR could exert a dual impact effect combining cytotoxic AML clearance with Gal-9-mediated signaling interference. Notably, the scFv incorporated in our construct derives from M6903, an antagonistic anti-TIM-3 antibody shown to block Gal-9 binding^29^, thereby providing a mechanistic basis for potential checkpoint pathway disruption by TIM-3.CAR-CIK cells. Based on this rationale, we investigated whether differential TIM-3 glycosylation could enable AML-selective CAR targeting.

Our biochemical analyses converge on a consistent model in which TIM-3 expressed on KASUMI-3 cells and primary AML blasts carries complex-type *N*-glycans that are hyper-fucosylated and hyper-sialylated. Strong AAL signal, its loss after PNGase F treatment, and the increased binding of a commercial anti–TIM-3 antibody (TIM-3-cmAb) following defucosylation (SGN-2FF) and desialylation (neuraminidase), collectively indicate extensive *N*-glycan elaboration, consistent with transcriptional upregulation of sialyl-and fucosyltransferases reported in AML^54,55,63–68^. A key finding is the partial PNGase F digestion seen in TIM-3 immunoprecipitations (IPs) (also with TIM-3scFv-mAb). Classical PNGase F resistance requires core α1,3-fucose, but in our case the persistence of HMW bands, still AAL-positive and sensitive to neuraminidase, is consistent with enrichment in sialylated Lewis structures (sLeˣ/sLeᵃ). These are *N*-glycan branches carrying α1,3/4-fucose on LacNAc together with α2,3-sialic acid. Their size and charge likely hinder full PNGase F access and create microheterogeneity, producing a persistent HMW smear^54,55^. This sLeˣ/ᵃ-high, AAL-high phenotype in blasts and KASUMI-3 cells matches AML glycomic and transcriptomic atlases^54,55,63–65,67,68^ and supports a glycoform-selective mechanism in which the TIM-3.CAR scFv binds efficiently to the heavily fucosylated and sialylated forms of TIM-3, while still retaining access to the polypeptide epitope after enzymatic trimming. These glycan features reflect broader metabolic rewiring in AML and may influence not only CAR engagement but also endogenous TIM-3 signaling. TIM-3/Gal-9 interaction, which is inherently glycan-dependent, promotes AML blasts survival and reinforces LSC self-renewal programs^20,60^. By targeting a glycoform-enriched TIM-3 population, our TIM-3.CAR may concomitantly eliminate LSCs and disrupt a pro-survival, immunosuppressive axis.

Overall, TIM-3 post-translational modifications remodel the accessibility of antibody epitopes. The TIM-3-cmAb used in our study exhibited weak binding to glycosylated TIM-3 on KASUMI-3 cells unless *N*-glycans were removed, indicating that AML-specific glycans mask the TIM-3-cmAb epitope. Conversely, the TIM-3scFv-mAb recognizes both glycosylated and deglycosylated TIM-3, but shows enhanced avidity toward the heavily glycosylated TIM-3 forms present in AML. This glycoform-dependent enhancement explains the preferential killing of KASUMI-3 and AML blasts and the low off-tumor reactivity toward healthy TIM-3⁺ immune cells. Indeed, from a safety perspective, our results indicate that monocytes and CIK cells present lower-complexity *N*-glycans, which might be insufficient to support productive CAR activation, creating a natural glycan-based safety window. Additionally, we show that inhibition of fucosylation using SGN-2FF abates TIM-3.CAR-CIK cytotoxicity and reduces effector-target avidity measured by z-Movi, further supporting the hypothesis that TIM-3 glycosylation acts not as an absolute requirement for TIM-3.CAR-mediated recognition, but as an avidity-boosting feature that selectively amplifies TIM-3.CAR binding to its protein-proximal epitope. In this context, glycans may act as structural stabilizers that enhance CAR binding avidity without constituting the primary recognition determinant—a concept recently illustrated by Park et al.^69^, who incorporated a non-signaling glyco-bridge binder targeting Tn-MUC1 into CAR-T cells to increase CAR–tumor cell avidity and cytotoxicity in pancreatic cancer models. While in that setting glycan engagement was engineered, in our context endogenous *N*-glycan remodeling on TIM-3 might serve as an intrinsic/endogenous glyco-bridge, amplifying CAR avidity and selectively reinforcing recognition of AML blasts.

Although TIM-3.CAR-CIK cells displayed potent cytotoxicity, cytokine release and proliferative activity against AML cell lines and primary AML blasts *in vitro*, AML intrinsic heterogeneity, including variable antigen expression levels among patients, represents a major limitation for monovalent CAR strategies. Consistent with this, TIM-3.CAR-CIK cells did not fully eradicate KASUMI-3 *in vivo*, likely due to heterogenous TIM-3 levels. To broaden the therapeutic window of this approach, we implemented an IF-BETTER Dual CAR platform. Specifically, we engineered next-generation Dual CAR-CIK cells co-targeting TIM-3 (using the previously characterized scFv) and CD33, a widely expressed and clinically validated myeloid marker^70–72^. These constructs (CD33.CAR/TIM-3.CCR and TIM-3.CAR/CD33.CCR) combine a second-generation CAR with a CCR, using CD28 and 4-1BB co-stimulatory domains *in trans*, a configuration supported by previous studies for enhancing CAR persistence and functionality^73–78^. Finally, we demonstrated that both Dual CARs can be efficiently expressed without evidence of fratricide.

Across AML models with variable antigen density, both configurations of Dual CAR-CIK cells showed robust, antigen-dependent activation and effective elimination of blasts and LSC-enriched subsets while sparing healthy TIM-3⁺ monocytes, NK cells, and CD34⁺ HSCs. These findings support IF-BETTER–mediated TIM-3/CD33 co-targeting as a strategy to broaden antigenic coverage while maintaining a favorable safety profile.

In conclusion, we identify AML-specific post-translational complex-type *N*-glycans on TIM-3, characterized by hyper-fucosylation and hyper-sialylation that potentiate TIM-3.CAR scFv activation and enable selective anti-leukemic activity. By integrating glycoform-boosted TIM-3 recognition with CD33 targeting through IF-BETTER logic gating, our strategy may improve tumor specificity, while limiting antigen escape and on-target/off-tumor toxicity. Moreover, TIM-3 targeting might interfere with immunosuppressive TIM-3/Gal-9 signaling and this dual impact effect warrants further investigations.

Together, these findings redefine antigen selectivity in AML and demonstrate that tumor-specific glycosylation patterns can be systematically leveraged to rationally design AML-restricted scFvs, with direct implications for the development of safer immunotherapies.

### Limitations of the study

This study links aberrant TIM-3 glycosylation and antigen accessibility to the enhanced AML-restricted activity of TIM-3.CAR and Dual CAR-CIK cells, although the specific contribution of distinct glycoforms to CAR engagement remains only partially elucidated. The mechanistic insights derived from *in vitro* biochemical assays may not fully capture the regulatory complexity present *in vivo*. Further studies will be required to clarify how variations in antigen density, glycan architecture, and features of the AML microenvironment influence the performance of TIM-3–directed strategies. Nonetheless, the present findings provide a solid framework for further optimization and future translational development.

## RESOURCE AVAILABILITY

### Lead contact

Requests for further information, resources, and reagents should be directed to and will be fulfilled by the lead contacts, Prof. Marta Serafini and Dr. Sarah Tettamanti

### Materials availability

Cell lines, the customized TIM-3 scFv–Fc antibody, CAR constructs, and relative sequences generated in this study are available upon request. Transfer of materials will require a material transfer agreement (MTA).

### Data and code availability

Raw source data generated and analyzed in this study are included in the published article and its supplementary information files. Analysis details are provided in the STAR Methods section. No standardized data types were generated in this study. This paper does not report original code. Any additional information required to reanalyze the data reported in this paper is available from the lead contact upon request.

## ACKNOWLEDGMENTS

This work was supported by grants from AIRC 5 × 1000 Immunity in Cancer Spreading and Metastasis (grant 21147), AIRC IG 2022 (grant 27507), AIRC IG 2018 (grant 22082), the Ministero della Salute Research project on CAR T cells for hematological malignancies and solid tumors conducted under the aegis of Alliance Against Cancer network, EU funding within the MUR PNRR “National Center for Gene Therapy and Drugs based on RNA Technology” (Project no. CN00000041 CN3 RNA), and MSCA grant MSCA_0000032, funded by the Italian Ministry of University and Research (MUR) through NextGeneration EU program.

The authors thank the parent committees Comitato Maria Letizia Verga, Quelli che…con Luca ODV, and Amici di Duccio for their generous and constant support. They also acknowledge Quelli che…con Luca ODV for financially supporting the PhD fellowship of C.A.

## AUTHOR CONTRIBUTIONS

Conceptualization, M.S. and S.T.; methodology, M.B., A.G., C.A., and S.T.; investigation, M.B., A.G., C.A, B.M., E.G., C.F., M.C.R., A.P., A.M.A., A.L. and B.C.; formal analysis, M.B., A.G., C.A. and B.M.; writing-original draft, M.B., A.G. and C.A., writing-review & editing, M.S., S.T., A.B. and G.G.; funding acquisition, M.S., G.G. and A.G.; supervision, M.S. and S.T. M.B., A.G., and C.A. contributed equally to this work. M.S. and S.T. jointly supervised this study.

## SUPPLEMENTAL INFORMATION

Supplemental File

## METHODS

### EXPERIMENTAL MODEL AND STUDY PARTICIPANT DETAILS

#### Cell lines

KASUMI-3, KG-1 (AML cell lines) and REH cell lines (ALL cell line) were sourced from American Type Culture Collection (ATCC). KASUMI-3 and KG-1 cells were maintained in RPMI 1640 medium (Sigma Aldrich) supplemented with 10% heat-inactivated fetal bovine serum (FBS, Gibco), 2mM L-glutamine, 25 IU/ml of penicillin and 25mg/ml of streptomycin (Lonza) at a concentration of 0.6×10^6^ cells/ml. The REH cell line was maintained in Advanced RPMI medium (Invitrogen) with 10% heat-inactivated FBS, 2mM L-glutamine, 25 IU/ml of penicillin and 25mg/ml of streptomycin, at a concentration of 1×10^6^ cells/ml. KG-1 TIM-3^+^ cells were generated by non-viral engineering of the parental wild-type KG-1 cell line.

The TIM-3 coding sequence was obtained from a commercially available Origene construct and subsequently cloned into a pT4 Sleeping Beauty transposon vector, which was then used to stably introduce TIM-3 into KG-1 cells. Briefly, 1×10^7^ cells were electroporated with the SF Cell Line 4D-Nucleofector^®^ X Kit L (Lonza) on 4D-Nucleofector^TM^ system (Lonza) using 7.5ug of supercoiled DNA transposon plasmid carrying the human TIM-3 sequence, and 2.5ug of supercoiled DNA encoding the SB100x transposase, following the manufacturer’s optimized protocol for KG-1 cells. TIM-3^+^ high cells were enriched by two rounds of sorting on a FACSAria cell sorter (BD Biosciences). The resulting KG-1 TIM-3^+^ cell line was maintained in RMPI 1640 with 10% heat-inactivated FBS, 2mM L-glutamine, 25 IU/ml of penicillin and 25mg/ml of streptomycin, at a concentration of 0.6×10^6^ cells/ml.

HMBS hTERT^+^ cells, an immortalized human bone marrow mesenchymal stromal cell line, was kindly provided by Dr. Campana (Department of Pediatrics, Yong Loo Lin School of Medicine, National University of Singapore). The HMBS hTERT^+^ cell line was maintained in RPMI 1640 supplemented with 10% heat-inactivated FBS, 2mM L-glutamine, 25 IU/ml of penicillin and 25mg/ml of streptomycin. All cell lines were periodically tested for mycoplasma contamination using a mycoplasma detection kit (Vazyme).

#### Mice

The study was approved by the Italian Ministry of Health. Experimental procedures involving animals were approved by the Institutional Animal Care and Use Committee (IACUC) at the Milano-Bicocca University (animal protocol ID 19/2022 PR – FB7CC.67.EXT.70) and performed in compliance with national and international laws and policies. NSG (NOD.Cg-*Prkdc^scid^ Il2rg^tm1Wjl^*/SzJ) mice were initially obtained from Charles River Laboratories and subsequently bred at the animal facility at the Milano-Bicocca University. Six- to eight-week-old male NSG mice were used for *in vivo* experiments and maintained in pathogen-free conditions. Schematics of the mouse models employed are provided in the main text. Mice were euthanized either at the end of the study or upon reaching pre-established IACUC health endpoints.

#### Human samples

Primary cytokine-induced killer cells (CIK) were generated from human peripheral blood mononuclear cells (PBMCs), obtained from healthy donors upon informed consent. Donor sex and age were not disclosed for privacy reasons. Bone marrow and peripheral blood samples were collected at diagnosis from adult (n= 41) and pediatric (n = 17) AML patients presenting different molecular and genetic subtypes. These samples were used to assess TIM-3 and CD33 expression by flow cytometry. Of 58 patients screened, 19 patients were female and 39 were male. Ten patient-derived specimens were further employed to evaluate the effector functions of TIM-3.CAR or Dual CAR-CIK cells. The study was approved by the Institutional Review Board of the Ethical Committee of Fondazione IRCCS San Gerardo dei Tintori Hospital (code: CAR 2018, MetaboCAR), and written informed consent was obtained from all patients in accordance with institutional guidelines and the Declaration of Helsinki.

## METHOD DETAILS

### Generation of CAR constructs

The high-affinity fully human scFv for the TIM-3 antigen was generated starting from the VH and VL sequence derived from an anti-TIM-3 mAb (M6903). The anti-TIM-3 scFv was cloned in frame with CH3-CD28-OX40-ζ, thus generating a TIM-3 specific third-generation CAR construct. The high-affinity, humanized rat anti-human CD33 scFv was generated using UCB’s Selected Lymphocyte Antibody Method (kindly provided by Dr. Helene Finney, UCB Celltech, Slough, 100 UK) and was cloned in frame with CH3-CD28-OX40-ζ, thus generating a CD33 specific third-generation CAR construct. Two different Dual CARs were generated (CD33.CAR/TIM-3.CCR and TIM-3.CAR/CD33.CCR) using scFvs from CD33.CAR and TIM-3.CAR. Dual CARs were characterized by a second-generation CAR (scFv cloned in frame with CH3 spacer and CD28 co-stimulatory domain), coupled with a chimeric costimulatory receptor (CCR) (scFv cloned in frame with CD8 stalk spacer and 4-1BB co-stimulatory domain). The CAR and CCR sequences forming each Dual CAR were linked by 2A peptide, codon optimized and cloned into an SB pT4-transposon vector. The SB-pT4 vector and SB100X transposase were kindly provided by Dr. Izsvak (Max-Delbruck-Center for Molecular Medicine, Berlin, Germany).

### CIK cell differentiation and engineering

CIK cells were generated, as previously described^30^, starting from human peripheral blood mononuclear cells (PBMCs) obtained from healthy donors. Briefly, PBMCs were isolated by Ficoll-Paque (Cytiva) density-gradient centrifugation. On day 0, PBMCs were plated at 3×10^6^ cells/ml in Advanced RPMI medium (supplemented with 10% FBS and 2mM L-glutamine, 25 IU/ml of penicillin and 25mg/ml of streptomycin) and interferon gamma (IFN-γ; 1000 U/mL; Dompe Biotec) was added. On day 1, interleukin-2 (IL-2; 300 U/mL;, Chiron BV) and anti-CD3 monoclonal antibody OKT3 (50 ng/ mL; Janssen-Cilag) were added. Electroporation was performed 24 hours later. For non-viral CAR engineering, 1×10^7^ CIK cells were resuspended in 100μl of Amaxa^TM^ nucleofector solution (P3 Primary Cell 4D-Nucleofector kit; Lonza) and electroporated using the 4D-Nucleofector^TM^ (Lonza) and the EO-115 program with 15μg supercoiled DNA transposon plasmid coding the CAR construct, and 0.5μg supercoiled DNA coding the SB100x transposase. Subsequently, fresh Advanced RPMI medium (supplemented as above) and IL-2 were added twice a week for 21 days of culture. At the end of the differentiation protocol, mature CIK cells consist predominantly of CD3^+^ T lymphocytes expressing CD8 and CD56 and exhibit an effector memory phenotype. Cell concentration was maintained at 1×10^6^ cells/ml throughout expansion.

### Flow cytometry

CIK cells were stained with fluorochrome-conjugated antibodies directed against CD3 (clone SK7), CD8 (clone RPA-T8), CD4 (clone SK3) (BioLegend), CD56 (clone B159), CD62L (clone DREG-56), CD45RO (clone UCHL1), and TIM-3 (clone 7D3) (BD Biosciences).

CIK cells exhaustion profile was also assessed by flow cytometry staining with antibodies directed against CD3 (clone SK7), PD-1 (clone EH12.1), and TIM-3 (clone 7D3), (BD Biosciences).

To establish CAR expression level, a TIM-3 fusion protein (Bio-techne) linked to the Fc portion of a human IgG_1_ and a recombinant human sialic acid binding Ig-Like Lectin 3/Siglec-3/CD33 protein with a Fc and a 6xHis tag at the C-terminus (C-Fc-6His) (Gentaur) were used, before proceeding with secondary staining with an anti-human IgG Fc Alexa Fluor 647-conjugated mAb (LiStarFish) and a 6xHis Tag (AD1.1.10) FITC-conjugated mAb (Invitrogen), respectively.

Leukemia cell lines were stained with TIM-3 (clone 7D3) (BD Biosciences) or TIM-3 (clone 344823) (R&D Systems) mAbs and CD33 (clone HIM3-4) (BD Biosciences), to verify the expression of the CAR target antigens. In *in vitro* assays, target cells were labeled with FITC- and PE-Cell Tracker (Invitrogen) or CellTrace^TM^ 155 Violet (Thermofisher Scientific) to distinguish them from effector cells. Cell death and apoptosis were detected using the Annexin V-Cy5 apoptosis detection kit (Enzo Life Sciences) or APC Annexin V Apoptosis Detection Kit (Biolegend) with 7-AAD according to the datasheet. Anti-human IFN-γ (clone B27), IL-2 (clone MQ1-17H129) and Ki67 (clone B56) mAbs (BD Biosciences) have been employed to evaluate CAR-CIK cells cytokine production and proliferation.

The Quantibrite^TM^ PE fluorescence quantitation kit (BD Biosciences), which enables the conversion of cell fluorescence intensity values into absolute numbers of receptors per cells, was employed to establish TIM-3 molecules/cell in KASUMI-3, AML blasts and healthy immune cells.

Human grafts in mice were analyzed using anti-human CD45 (clone 2D1), CD3, TIM-3, CD33 (clone P67.6) (BD Biosciences) and anti-mouse CD45 (clone 30-F11) mAbs (ThermoFisher). The absolute number of CAR-CIK and leukemic cells was quantified by flow cytometry, after adding 10μl of CountBright^TM^ Absolute Counting Beads (Thermo Fisher Scientific) per tube. Samples were analyzed using a FACS CANTO II flow cytometer and LSR Fortessa cell analyzer (BD Biosciences). All data were analyzed with the FlowJo software (Tree Star Inc.).

### Short-term and long-term cytotoxicity assays

In the short-term cytotoxicity assay, CIK cells were seeded to a 96-well round-bottom plate for 4 hours at an Effector:Target (E:T) ratio of 5:1 (1×10^5^ effector cells) with the target cells (previously labeled with FITC-cell tracker), KASUMI-3, KG-1 and KG1 TIM-3^+^ or with REH, a CD33^-^ TIM-3^-^ ALL cell line used as a negative control. Target cell killing was measured through apoptosis detection as previously described. The percentage of killed target cells was calculated as follows:

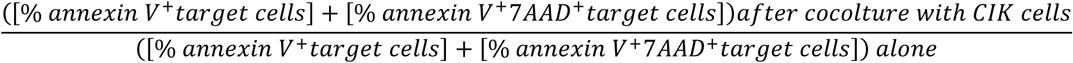

In the long-term cytotoxic assay, CIK cells were co-cultured in 96-well round-bottom plate for 7 days at different E:T ratios (1:10, 1:50, 1:100) (2×10^4^ target cells) against target AML cell lines (pre-labeled with FITC or CellTrace Violet) or against primary AML samples (pre-labeled with PE or CellTrace Violet), which were previously seeded on a layer of immortalized human bone marrow mesenchymal stromal cells (HBMS hTERT^+^). The percentage of target cells survival was calculated by flow cytometry, after adding 10μl of CountBright^TM^ Absolute Counting Beads per tube, using the following formula:

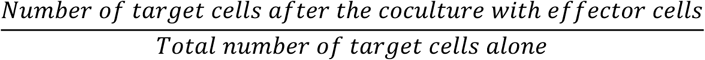

Data were analyzed using Flowjo and graphs were generated in GraphPad Prism.

### Proliferation assay

*In vitro* proliferation assays were performed in 96-well round-bottom plates. CAR-CIK cells were co-cultured with irradiated cell lines (KASUMI-3, KG-1, REH) or primary AML patient samples at an E:T ratio of 1:1 (1×10^5^ effectors). To evaluate the specificity of CAR-CIK cell proliferative response, the ALL cell line REH was employed as negative control. After a 72 hours co-culture, cells were collected and stained for CD3 and CAR expression, and CAR-CIK proliferation ability was evaluated by Ki67 (clone B56, BD Biosciences) intracellular staining by flow cytometry. FlowJo was used for data analysis, while GraphPad Prism was used to generate the graphs.

### Intracellular cytokine staining (ICS)

Cytokine-producing ability of CAR-CIK cells was evaluated after a 5-hour co-culture with the different target cells and primary AML samples at an E:T ratio of 1:3 (1×10^5^ effectors) in a 96-well round-bottom plate. After 2.5 hours, BD GolgiStop^TM^ was added. The co-culture was then maintained for additional 2.5 hours, after which cells were collected and stained for CD3 and CAR expression. Intracellular cytokine staining for IL-2 (clone MQ1-17H129, BD Biosciences) and IFN-γ (clone B27, BD Biosciences) was performed using the BD Cytofix/Cytoperm kit, according to the datasheet. Samples were then acquired by flow cytometry. Data analysis was performed with FlowJo, and graphs were created in GraphPad Prism.

### ELISA

KASUMI-3, AML blasts and REH cells (2×10^6^ cells) were cultured in RPMI 1640 medium 10% FBS, 2mM L-glutamine, 25 IU/ml of penicillin and 25mg/ml of streptomycin for 24 hours, before supernatant harvesting. The levels of Gal-9 were assessed in culture supernatants using a commercially available ELISA kit (Human Galectin-9, DuoSet ELISA, R&D Systems), according to the manufacturer’s instructions. Absorbance readings were obtained using the Spark® Multimode Microplate Reader (TECAN).

### Monocytes and NK-cells sorting

Monocytes were enriched from PBMCs using human CD14 MicroBeads (Miltenyi Biotec) according to the manufacturer’s instructions. After CD14^+^ cells sorting, the negative fraction obtained from monocytes selection was itself sorted adopting CD16 MicroBeads (Miltenyi Biotec), thus obtaining a CD14^-^ CD16^+^ NK cell enriched population. Monocytes and NK cells purification was evaluated by flow cytometry, using CD14 (clone M5E2) (BD Biosciences) and CD16 (NKP15) (BD Bioscience). TIM-3 and CD33 expression on monocytes and NK cells was assessed by flow cytometry, employing CD33 (P67.6) (BD Bioscience), TIM-3 (clone 7D3) (BD Biosciences) and TIM-3 (clone 344823) (R&D Systems) antibodies.

### Analysis of TIM-3–directed fratricide and on-target/off-tumor toxicity

To assess if TIM-3.CAR-CIK cells may undergo self-killing or mediate a cytotoxic effect on activated TIM-3^+^ unmodified CIK cells, fratricide co-culture assays were performed at an E:T ratio of 5:1 (1×10^5^ effectors). TIM-3^+^ unmanipulated CIK cells, expanded from healthy donors, were labeled with FITC cell tracker and seeded. TIM-3.CAR-CIK cells were then added in the corresponding wells. After 4-hour incubation, target cell killing was measured through apoptosis detection by flow cytometry, via Annexin V and 7-AAD staining. The percentage of killed target cells was calculated as previously reported.

To evaluate the potential TIM-3.CAR-, CD33.CAR/TIM3.CCR- and TIM3.CAR/CD33.CCR-CIK cells cytotoxicity against TIM-3^+^ monocytes, unmodified CIK cell and NK cell subsets, *in vitro* killing assays were performed. CAR-CIK cells and TIM-3^+^ target cells were co-cultured at an E:T ratio of 5:1 (1×10^5^ effectors). Target cells were previously stained with FITC cell-tracker. After a 4 hours incubation, target cell killing was established by flow cytometry, after adding 10μl of CountBright^TM^ Absolute Counting Beads (Thermo Fisher Scientifics) per tube with the previously reported formula. All assays were performed in 96-well round-bottom plates. Data analysis was performed with FlowJo, and graphs were created in GraphPad Prism.

### Protein extraction and BCA quantification

KASUMI-3, monocytes, NT were lysed in RIPA lysis buffer containing (1% of NP-40, 0.5% Na-Deoxyxholate, 350mM NaCl, 0.1% SDS; Sigma Aldrich) supplemented of 1% of Protease Inhibitor Cocktail (SigmaAldrich) and 0.25mM of Phenyl-methyl-sulfonyl fluoride (Sigma Aldrich). After 30 minutes of ice incubation, the samples were centrifuged at +4°C for 10 minutes at 21.000*g* to remove cell debris. Supernatant was retrieved, and stocked at −80° until usage. Protein quantification was performed with Bicinchoninic Acid Assay (BCA-Thermo-Scientific™ Pierce™) according to manufacturer Microplate procedure protocol. The standard curve was obtained with 8 serial dilutions of BSA protein, included in the kit and the assay was performed in duplicate. 10ul of standard curve or protein sample were combined with 200ul of Working Reagent and incubated at 37°C for 20 minutes. The absorbance at 562nm was detected with Spark^®^ Multimode Microplate Reader (TECAN).

### Immunoprecipitation

Immunoprecipitation of TIM-3 from total extract lysate of KASUMI-3, monocytes and T cells was performed following A/G Beads (Thermo-Scientific™ Pierce™) protocol. Specifically, 5μg of Human TIM-3 mAb (Clone 344801) (Biotechne) were combined with 300μg of KASUMI-3 or 150μg of monocytes and T cells total lysate and the reaction volume adjust to 500μl with RIPA. The reaction was incubated over-night at +4°C with mixing. The following day, 25μl of Pierce Protein A/G Magnetic Beads (Thermo-Scientific™ Pierce™) were primed and washed twice with Wash Buffer. The antibody/sample reactions were added to the beads and incubated for 1 hour at room temperature with mixing. Beads-antibody-sample complexes were collected with a magnetic stand and the flowthrough stored for the analysis. The complexes were washed twice with Wash Buffer and incubated with 100μl of Low-pH-Eluition Buffer for 10 minutes at room temperature with mixing. The beads were separated with the magnetic field and the supernatant, containing the target protein and antibody, neutralized with 15μl of Neutralization Buffer. Total amount of TIM-3 protein retrieved from each cell population is quantified with BCA assay and TIM-3 enrichment was validated with Western Blot analysis.

### Deglycosylation (PNGase F, Neuraminidase A, O-glycosidases) and immunoblot/lectin blot

All experiments were performed on total cellular protein extracts (whole-cell lysates); where indicated, immunoprecipitated (IP) eluates were also analyzed. For immunoprecipitation-based immunoblots, TGX (Tris-Glycine eXtended) stain-free total protein signal was used as a total protein loading/normalization control. PNGase F, Neuraminidase A, and O-glycosidases treatments were carried out according to the manufacturer’s instructions (New England BioLabs; PNGase F kit P0704S; Neuraminidase A kit P0722S; O-Glycosidase, endo-α-N-acetylgalactosaminidase NEB P0733).

Briefly, 0.2 µg protein from lysates or IP eluates were denatured in the supplied buffer (0.5% SDS, 40 mM DTT) for 10 min at 100 °C. After cooling, the appropriate GlycoBuffer was added per protocol. For *N-*glycan removal, samples were incubated with PNGase F at 37 °C for 1 h. For O-glycan removal, samples were first desialylated with Neuraminidase A, then incubated with O-Glycosidase (± accessory exoglycosidases as above) at 37 °C for 1 h. Control samples were processed identically without adding deglycosylating enzymes.

Proteins were resolved by SDS–PAGE on 12% Bis–Tris precast gels (Bio-Rad) or, for some experiments, on 4–20% gradient gels (Bio-Rad) and transferred to PVDF membranes using the Trans-Blot Turbo system (Bio-Rad) following the manufacturer’s instructions. Membranes were probed with anti-human TIM-3 antibody (TIM-3-cmAb) (1:250; R&D Systems, MAB23652), a recombinant monoclonal antibody derived from the TIM-3.CAR scFv (TIM-3 scFv-mAb) (1:500; GENEWIZ), biotinylated Aleuria aurantia lectin (AAL; 1:3000; Vector Laboratories, B-1395-1), and biotinylated Ricinus communis agglutinin I (RCA I; 1:3000; Vector Laboratories, B-1085-1). Where indicated, membranes were additionally probed with anti–sialyl-Lewis X (CSLEX1 Biosciences Cat# 551344) and anti–sialyl-Lewis A (CA19-9 ThermoFisher Cat# MA5-42322) antibodies (each 1:500) to detect sialyl-Lewis–type determinants on immunoprecipitated TIM-3. Primary antibodies and horseradish peroxidase–conjugated secondary reagents were diluted in 5% (w/v) non-fat milk in TBST. Signals were developed with ECL Western Blotting detection reagent (Bio-Rad) and imaged on a ChemiDoc MP system (Bio-Rad).

### Assessment of TIM-3.CAR-CIK cytotoxicity against SGN-2FF–treated KASUMI-3

KASUMI-3 cells (2×10^5^) were seeded in 24-well plates and either left untreated or exposed to fucosyltransferase inhibitor, 2F-Peracetyl-Fucose (SGN-2FF) (Sigma Aldrich) (100nM, 48 hours). After treatment, the ability of TIM-3.CAR-CIK cell to recognize and eliminate SGN-2FF-treated or control KASUMI-3 was assessed in co-culture using a range of E:T ratios (5:1, 1:1, 0.5:1, 0.25:1 and 0.125:1). Cytotoxicity assays were performed in 96 well round-bottom plates using 1×10^5^ effector cells per condition. The percentage of killed target cells was calculated as previously reported.

### Cell avidity measurement using z-Movi

TIM-3.CAR-CIK cell binding avidity against SGN-2FF-treated and control KASUMI-3 monolayers was measured using a z-Movi Cell Avidity Analyzer (LUMICKS). Briefly, z-Movi chips were coated with poly-L-lysine prior to attaching a monolayer of leukemic cells. 2,5×10^6^ KASUMI-3 cells were seeded on each z-Movi chip at a concentration of 1,25×10^8^ cells/mL and cultured at 37°C in a dry incubator, 2 hours and a half in advance. TIM-3.CAR-CIK cells (2×10^5^ CAR-CIK cells stained with Cell Tracker Deep Red Dye) (ThermoFisher) were serially flowed in the chips and incubated on the SGN-2FF treated or control KASUMI-3 monolayer for 5 min before initializing the protocol standard linear force ramp. Data are presented as mean % bound T cells throughout the acquisition and at the maximum mean force applied, 1000 piconewtons, pN. This was repeated for six biological CIK cell donors. Avidity experiments were conducted according to the manufacturer (LUMICKS) instructions and recommendations, and cell avidity was quantified using the z-Movi software (v1.0).

### Quantitative real-time PCR

Total RNA was extracted from human cells (KASUMI-3, primary AML blasts and monocytes) with QIAZOL reagent (Qiagen) using the miRNeasy Micro Kit (Qiagen) following the manufacturer’s instructions. 1μg of total RNA was reversed transcribed to synthesize cDNA, using SuperScript™ II Reverse Transcriptase (ThermoFisher) in the presence of random hexamers (Invitrogen). Real-time PCR was performed in duplicate in the QuantStudio™ 3 Real-Time PCR System (ThermoFisher), using SYBR Green PCR Master Mix (ThermoFisher) and SYBR Green probes. SYBR Green primers for the following genes were purchased from metabion or macrogen: 18S, FUT7, FUT8, ST3GAL3, ST3GAL4 and ST3GAL6. Data were normalized to 18S RNA gene expression. The relative mRNA expression was calculated by the comparative threshold cycle (Ct) method. The results are expressed as fold change as indicated in the graphs.

### Xenograft animal models

For *in vivo* experiments, six- to eight-week-old male NSG mice were sublethally irradiated (200 cGγ) and intravenously infused with TIM-3^+^ KASUMI-3 cell line (10^7^ cells per mouse in 100μl PBS). Two weeks later, mice received weekly tail vein injections of 10^7^ TIM-3.CAR^+^-CIK cells for 3 consecutive weeks or were left untreated. Animals were euthanized one week after the last infusion, and peripheral blood (PB), spleen and bone marrow (BM) were analyzed by flow cytometry to establish the residual AML burden.

For *in vivo* evaluation of Dual CAR-CIK cell activity, low-burden and high-burden AML xenograft models were generated by transplanting NSG mice with KG-1 TIM-3^+^ (CD33^high^TIM-3^high^) AML cell line (5×10^6^ cells per mouse in 100μl PBS). In the low-burden setting, mice received CAR-CIK infusions on day 5 and 12 after leukemia inoculation. In the high burden model, CAR-CIK treatment was administered to NSG mice at day 14 and day 21 after KG-1 TIM-3^+^ inoculation. CAR-CIK cells were delivered by tail-vein injection (10^7^ cells per mouse in 100μl PBS). At endpoint, PB, spleen and BM were retrieved to assess CAR-CIK cell persistence and leukemia elimination.

Spleen and BM were collected and processed into single-cell suspensions. Briefly, after harvesting spleens were gently mashed using the flat end of a syringe plunger, while BM was isolated by centrifugation at 13.000 rpm for 5 min. PB was collected from NSG mice via cardiac puncture at sacrifice. All samples were lysed with ACK lysis buffer (Voden Medical) for subsequent flow cytometry staining and analysis. Flow cytometry data were processed using FlowJo, while graphs were generated with GraphPad Prism.

### CFU assay

PBMCs were isolated from cord blood by Ficoll-Paque separation and CD34^+^ cells were purified using human CD34 MicroBead Kit (Miltenyi Biotec) following the manufacturer’s instructions. Purity was assessed by flow cytometry, with CD34 Ab (clone 8G12) (BD Bioscience) staining. A total of 5×10^3^ CD34^+^ cells per condition were co-cultured with NT, CD33.CAR-, CD33.CAR/TIM-3.CCR-, TIM-3.CAR-, TIM-3.CAR/CD33.CCR-CIK cells with E:T ratio 10:1 (5×10⁴ effector cells) in Advanced RPMI medium (supplemented with 10% FBS and 2mM L-glutamine, 25 IU/ml of penicillin and 25mg/ml of streptomycin) at 37°C and 5% CO₂. After 24-hours, 50μl of each condition of the co-culture were mixed with 2.5ml of MethoCult (H4434 Classic) (Voden Medical) and plated into 6-well plates following the manufacturer’s instructions. Colony formation was assessed microscopically after 14-16 days; total CFU counts were recorded for each condition.

## QUANTIFICATION AND STATISTICAL ANALYSIS

Data were analyzed using GraphPad Prism 10 Software version (GraphPad Software Inc, La Jolla, CA, USA). All data are presented as mean +/- SD. Two-tailed unpaired or paired Student’s t-test between the two groups was used to determine significance. Two-way ANOVA was performed for grouped statistics. Differences with a p value < 0.05 were considered statistically significant (* p< 0.05; ** p< 0.01; *** p< 0.001; ***** p< 0.0001). One-way ANOVA was performed for grouped statistics and differences with a p value < 0,05 were considered statistically significant (* p< 0.05; ** p< 0.01; *** p< 0.001; ***** p< 0.0001). All *in vitro* data presented are representative of at least three biological replicates. *In vivo* animal studies were performed using at least two biological replicates. Illustrations were created by BioRender.com.

## REFERENCES

1. Shimony, S., Stahl, M., and Stone, R.M. (2023). Acute myeloid leukemia: 2023 update on diagnosis, risk-stratification, and management. Am. J. Hematol. 98, 502–526. 10.1002/AJH.26822.

2. Boscaro, E., Urbino, I., Catania, F.M., Arrigo, G., Secreto, C., Olivi, M., D’Ardia, S., Frairia, C., Giai, V., Freilone, R., et al. (2023). Modern Risk Stratification of Acute Myeloid Leukemia in 2023: Integrating Established and Emerging Prognostic Factors. Cancers (Basel). 15. 10.3390/CANCERS15133512.

3. Döhner, H., Wei, A.H., Appelbaum, F.R., Craddock, C., DiNardo, C.D., Dombret, H., Ebert, B.L., Fenaux, P., Godley, L.A., Hasserjian, R.P., et al. (2022). Diagnosis and management of AML in adults: 2022 recommendations from an international expert panel on behalf of the ELN. Blood 140, 1345–1377. 10.1182/BLOOD.2022016867.

4. Ishikawa, F., Yoshida, S., Saito, Y., Hijikata, A., Kitamura, H., Tanaka, S., Nakamura, R., Tanaka, T., Tomiyama, H., Saito, N., et al. (2007). Chemotherapy-resistant human AML stem cells home to and engraft within the bone-marrow endosteal region. Nat. Biotechnol. 2007 2511 25, 1315–1321. 10.1038/nbt1350.

5. Döhner, H., Weisdorf, D.J., and Bloomfield, C.D. (2015). Acute Myeloid Leukemia. N. Engl. J. Med. 373, 1136–1152. 10.1056/NEJMra1406184.

6. Kreidieh, F., Abou Dalle, I., Moukalled, N., El-Cheikh, J., Brissot, E., Mohty, M., and Bazarbachi, A. (2022). Relapse after allogeneic hematopoietic stem cell transplantation in acute myeloid leukemia: an overview of prevention and treatment. Int. J. Hematol. 116, 330–340. 10.1007/S12185-022-03416-7.

7. Rampotas, A., and Roddie, C. (2025). The present and future of CAR T-cell therapy for adult B-cell ALL. Blood 145, 1485–1497. 10.1182/BLOOD.2023022922.

8. Zhang, X., Zhu, L., Zhang, H., Chen, S., and Xiao, Y. (2022). CAR-T Cell Therapy in Hematological Malignancies: Current Opportunities and Challenges. Front. Immunol. 13, 1–20. 10.3389/fimmu.2022.927153.

9. Saito, S., and Nakazawa, Y. (2024). CAR-T cell therapy in AML: recent progress and future perspectives. Int. J. Hematol. 120, 455–466. 10.1007/S12185-024-03809-W.

10. Tettamanti, S., Pievani, A., Biondi, A., Dotti, G., and Serafini, M. (2022). Catch me if you can: how AML and its niche escape immunotherapy. Leukemia 36, 13–22. 10.1038/S41375-021-01350-X.

11. Atilla, E., and Benabdellah, K. (2023). The Black Hole: CAR T Cell Therapy in AML. Cancers (Basel). 15, 2713. 10.3390/CANCERS15102713.

12. Khalifeh, M., Hopewell, E., and Salman, H. (2025). CAR-T cell therapy for treatment of acute myeloid leukemia, advances and outcomes. Mol. Ther. 33, 2441–2453. 10.1016/J.YMTHE.2025.03.052.

13. Liu, Y., Wang, W., Wang, C., Deng, J., Hu, Y., Mei, H., and Luo, S. (2025). Recent advances of chimeric antigen receptor T-cell therapy for acute myeloid leukemia. Front. Immunol. 16, 1572407. 10.3389/FIMMU.2025.1572407/XML.

14. Kikushige, Y., Shima, T., Takayanagi, S.I., Urata, S., Miyamoto, T., Iwasaki, H., Takenaka, K., Teshima, T., Tanaka, T., Inagaki, Y., et al. (2010). TIM-3 is a promising target to selectively kill acute myeloid leukemia stem cells. Cell Stem Cell 7, 708–717. 10.1016/j.stem.2010.11.014.

15. Kikushige, Y., and Akashi, K. (2012). TIM-3 as a therapeutic target for malignant stem cells in acute myelogenous leukemia. Ann. N. Y. Acad. Sci. 1266, 118–123. 10.1111/j.1749-6632.2012.06550.x.

16. Kikushige, Y., and Miyamoto, T. (2015). Identification of TIM-3 as a Leukemic Stem Cell Surface Molecule in Primary Acute Myeloid Leukemia. Oncology 89, 28–32. 10.1159/000431062.

17. Wang, Z., Chen, J., Wang, M., Zhang, L., and Yu, L. (2021). One Stone, Two Birds: The Roles of Tim-3 in Acute Myeloid Leukemia. Front. Immunol. 12. 10.3389/FIMMU.2021.618710.

18. Darwish, N.H.E., Sudha, T., Godugu, K., Elbaz, O., Abdelghaffar, H.A., Hassan, E.E.A., and Mousa, S.A. (2016). Acute myeloid leukemia stem cell markers in prognosis and targeted therapy: potential impact of BMI-1, TIM-3 and CLL-1. Oncotarget 7, 57811. 10.18632/ONCOTARGET.11063.

19. Jan, M., Chao, M.P., Cha, A.C., Alizadeh, A.A., Gentlese, A.J., Weissmana, I.L., and Majeti, R. (2011). Prospective separation of normal and leukemic stem cells based on differential expression of TIM3, a human acute myeloid leukemia stem cell marker. Proc. Natl. Acad. Sci. U. S. A. 108, 5009–5014. 10.1073/pnas.1100551108.

20. Kikushige, Y., Miyamoto, T., Yuda, J., Jabbarzadeh-Tabrizi, S., Shima, T., Takayanagi, S.I., Niiro, H., Yurino, A., Miyawaki, K., Takenaka, K., et al. (2015). A TIM-3/Gal-9 Autocrine Stimulatory Loop Drives Self-Renewal of Human Myeloid Leukemia Stem Cells and Leukemic Progression. Cell Stem Cell 17, 341–352. 10.1016/j.stem.2015.07.011.

21. Gonçalves Silva, I., Rüegg, L., Gibbs, B.F., Bardelli, M., Fruewirth, A., Varani, L., Berger, S.M., Fasler-Kan, E., and Sumbayev, V. V. (2016). The immune receptor Tim-3 acts as a trafficker in a Tim-3/galectin-9 autocrine loop in human myeloid leukemia cells. Oncoimmunology 5. 10.1080/2162402X.2016.1195535.

22. Gonçalves Silva, I., Yasinska, I.M., Sakhnevych, S.S., Fiedler, W., Wellbrock, J., Bardelli, M., Varani, L., Hussain, R., Siligardi, G., Ceccone, G., et al. (2017). The Tim-3-galectin-9 Secretory Pathway is Involved in the Immune Escape of Human Acute Myeloid Leukemia Cells. EBioMedicine 22, 44–57. 10.1016/J.EBIOM.2017.07.018.

23. Mao, C., Li, J., Feng, L., and Gao, W. (2023). Beyond antibody fucosylation: α-(1,6)-fucosyltransferase (Fut8) as a potential new therapeutic target for cancer immunotherapy. Antib. Ther. 6, 87–96. 10.1093/ABT/TBAD004.

24. Liu, B., Ma, H., Liu, Q., Xiao, Y., Pan, S., Zhou, H., and Jia, L. (2019). MiR-29b/Sp1/FUT4 axis modulates the malignancy of leukemia stem cells by regulating fucosylation via Wnt/β-catenin pathway in acute myeloid leukemia. J. Exp. Clin. Cancer Res. 38. 10.1186/S13046-019-1179-Y.

25. Filipsky, F., and Läubli, H. (2024). Regulation of sialic acid metabolism in cancer. Carbohydr. Res. 539, 109123. 10.1016/J.CARRES.2024.109123.

26. Stanczak, M.A., and Läubli, H. (2023). Siglec receptors as new immune checkpoints in cancer. Mol. Aspects Med. 90. 10.1016/J.MAM.2022.101112.

27. Haubner, S., Perna, F., Köhnke, T., Schmidt, C., Berman, S., Augsberger, C., Schnorfeil, F.M., Krupka, C., Lichtenegger, F.S., Liu, X., et al. (2018). Coexpression profile of leukemic stem cell markers for combinatorial targeted therapy in AML. Leuk. 2018 331 33, 64–74. 10.1038/s41375-018-0180-3.

28. Willier, S., Rothämel, P., Hastreiter, M., Wilhelm, J., Stenger, D., Blaeschke, F., Rohlfs, M., Kaeuferle, T., Schmid, I., Albert, M.H., et al. (2021). CLEC12A and CD33 coexpression as a preferential target for pediatric AML combinatorial immunotherapy. Blood 137, 1037–1049. 10.1182/BLOOD.2020006921.

29. Zhang, D., Jiang, F., Zaynagetdinov, R., Huang, H., Sood, V.D., Wang, H., Zhao, X., Jenkins, M.H., Ji, Q., Wang, Y., et al. (2020). Identification and characterization of M6903, an antagonistic anti–TIM-3 monoclonal antibody. Oncoimmunology 9. 10.1080/2162402X.2020.1744921.

30. Rotiroti, M.C., Buracchi, C., Arcangeli, S., Galimberti, S., Valsecchi, M.G., Perriello, V.M., Rasko, T., Alberti, G., Magnani, C.F., Cappuzzello, C., et al. (2020). Targeting CD33 in Chemoresistant AML Patient-Derived Xenografts by CAR-CIK Cells Modified with an Improved SB Transposon System. Mol. Ther. 28, 1974–1986. 10.1016/j.ymthe.2020.05.021.

31. Yang, R., Sun, L., Li, C.F., Wang, Y.H., Yao, J., Li, H., Yan, M., Chang, W.C., Hsu, J.M., Cha, J.H., et al. (2021). Galectin-9 interacts with PD-1 and TIM-3 to regulate T cell death and is a target for cancer immunotherapy. Nat. Commun. 12. 10.1038/s41467-021-21099-2.

32. Kenderian, S.S., Ruella, M., Shestova, O., Klichinsky, M., Kim, M., Porter, D.L., June, C.H., and Gill, S. (2016). Identification of PD1 and TIM3 As Checkpoints That Limit Chimeric Antigen Receptor T Cell Efficacy in Leukemia. Biol. Blood Marrow Transplant. 22, S19–S21. 10.1016/j.bbmt.2015.11.291.

33. Pabst, C., Bergeron, A., Lavallée, V.P., Yeh, J., Gendron, P., Norddahl, G.L., Krosl, J., Boivin, I., Deneault, E., Simard, J., et al. (2016). GPR56 identifies primary human acute myeloid leukemia cells with high repopulating potential in vivo. Blood 127, 2018–2027. 10.1182/BLOOD-2015-11-683649.

34. Reily, C., Stewart, T.J., Renfrow, M.B., and Novak, J. (2019). Glycosylation in health and disease. Nat. Rev. Nephrol. 15, 346–366. 10.1038/S41581-019-0129-4.

35. Posey, A.D., Schwab, R.D., Boesteanu, A.C., Steentoft, C., Mandel, U., Engels, B., Stone, J.D., Madsen, T.D., Schreiber, K., Haines, K.M., et al. (2016). Engineered CAR T Cells Targeting the Cancer-Associated Tn-Glycoform of the Membrane Mucin MUC1 Control Adenocarcinoma. Immunity 44, 1444–1454. 10.1016/J.IMMUNI.2016.05.014.

36. Miyoshi, E., Moriwaki, K., Terao, N., Tan, C.C., Terao, M., Nakagawa, T., Matsumoto, H., Shinzaki, S., and Kamada, Y. (2012). Fucosylation is a promising target for cancer diagnosis and therapy. Biomolecules 2, 34–45. 10.3390/BIOM2010034.

37. Rodrigues, E., and Macauley, M.S. (2018). Hypersialylation in Cancer: Modulation of Inflammation and Therapeutic Opportunities. Cancers (Basel). 10. 10.3390/CANCERS10060207.

38. Gorman, J. V., and Colgan, J.D. (2014). Regulation of T cell responses by the receptor molecule Tim-3. Immunol. Res. 59, 56–65. 10.1007/S12026-014-8524-1.

39. Chongsaritsinsuk, J., Steigmeyer, A.D., Mahoney, K.E., Rosenfeld, M.A., Lucas, T.M., Smith, C.M., Li, A., Ince, D., Kearns, F.L., Battison, A.S., et al. (2023). Glycoproteomic landscape and structural dynamics of TIM family immune checkpoints enabled by mucinase SmE. Nat. Commun. 14. 10.1038/S41467-023-41756-Y.

40. Sanmartín-Martínez, J., Wiersma, V.R., and Marneth, A.E. (2024). The sweet symphony of N-glycans in myeloid malignancies. Front. Hematol. 3, 1–13. 10.3389/frhem.2024.1415618.

41. Hegde, R., and Podder, S.K. (1998). Evolution of tetrameric lectin Ricinus communis agglutinin from two variant groups of ricin toxin dimers. Eur. J. Biochem. 254, 596–601. 10.1046/J.1432-1327.1998.2540596.X.

42. Wu, A.M., Wu, J.H., Singh, T., Lai, L.J., Yang, Z., and Herp, A. (2006). Recognition factors of Ricinus communis agglutinin 1 (RCA(1)). Mol. Immunol. 43, 1700–1715. 10.1016/J.MOLIMM.2005.09.008.

43. Pizzitola, I., Anjos-Afonso, F., Rouault-Pierre, K., Lassailly, F., Tettamanti, S., Spinelli, O., Biondi, A., Biagi, E., and Bonnet, D. (2014). Chimeric antigen receptors against CD33/CD123 antigens efficiently target primary acute myeloid leukemia cells in vivo. Leuk. 2014 288 28, 1596–1605. 10.1038/leu.2014.62.

44. Kim, M.Y., Yu, K.-R., Kenderian, S.S., Ruella, M., Chen, S., Shin, T.-H., Aljanahi, A.A., Schreeder, D., Klichinsky, M., Shestova, O., et al. (2018). Genetic Inactivation of CD33 in Hematopoietic Stem Cells to Enable CAR T Cell Immunotherapy for Acute Myeloid Leukemia. Cell 173, 1439. 10.1016/J.CELL.2018.05.013.

45. Sakoda, T., Kikushige, Y., Miyamoto, T., Irifune, H., Harada, T., Hatakeyama, K., Kunisaki, Y., Kato, K., and Akashi, K. (2023). TIM-3 signaling hijacks the canonical Wnt/β-catenin pathway to maintain cancer stemness in acute myeloid leukemia. Blood Adv. 7, 2053–2065. 10.1182/BLOODADVANCES.2022008405.

46. Zhang, X., Feng, Z., Pranatharthi Haran, A., and Hua, X. (2025). Dual nanobody-redirected and Bi-specific CD13/TIM3 CAR T cells eliminate AML xenografts without toxicity to human HSCs. Oncoimmunology 14. 10.1080/2162402X.2025.2458843.

47. Lee, W.H.S., Ye, Z., Cheung, A.M.S., Goh, Y.P.S., Oh, H.L.J., Rajarethinam, R., Yeo, S.P., Soh, M.K., Chan, E.H.L., Tan, L.K., et al. (2021). Effective Killing of Acute Myeloid Leukemia by TIM-3 Targeted Chimeric Antigen Receptor T Cells. Mol. Cancer Ther. 20, 1702–1712. 10.1158/1535-7163.MCT-20-0155.

48. Klaihmon, P., Luanpitpong, S., Kang, X., and Issaragrisil, S. (2023). Anti-TIM3 chimeric antigen receptor-natural killer cells from engineered induced pluripotent stem cells effectively target acute myeloid leukemia cells. Cancer Cell Int. 23, 297-. 10.1186/S12935-023-03153-9/FIGURES/7.

49. Pe, K.C.S., Jewmoung, S., Rad, S.A.H., Chantarat, N., Chanswangphuwana, C., Tashiro, H., Suppipat, K., and Tawinwung, S. (2024). Optimization of anti-TIM3 chimeric antigen receptor with CD8α spacer and TNFR-based costimulation for enhanced efficacy in AML therapy. Biomed. Pharmacother. 179, 117388. 10.1016/J.BIOPHA.2024.117388.

50. Abrantes, R., Forcados, C., Warren, D.J., Santos-Ferreira, L., Fleten, K.G., Senra, E., Costa, A.F., Krpina, K., Henrique, R., Liberg, A.M., et al. (2025). Pan-carcinoma sialyl-Tn-targeting expands CAR therapy to solid tumors. Cell Reports Med. 6. 10.1016/j.xcrm.2025.102350.

51. Zingg, A., Ritschard, R., Thut, H., Buchi, M., Holbro, A., Oseledchyk, A., Heinzelmann, V., Buser, A., Binder, M., Zippelius, A., et al. (2025). Targeting Cancer-Associated Glycosylation for Adoptive T-cell Therapy of Solid Tumors. Cancer Immunol. Res. 13, 990–1003. 10.1158/2326-6066.CIR-24-1050.

52. Heinzelbecker, J., Fauskanger, M., Jonson, I., Krengel, U., Løset, G.Å., Munthe, L., and Tveita, A. (2024). Chimeric antigen receptor T cells targeting the GM3(Neu5Gc) ganglioside. Front. Immunol. 15, 1–12. 10.3389/fimmu.2024.1331345.

53. Jetani, H., Navarro-Bailón, A., Maucher, M., Frenz, S., Verbruggen, C., Yeguas, A., Vidriales, M.B., González, M., Rial Saborido, J., Kraus, S., et al. (2021). Siglec-6 is a novel target for CAR T-cell therapy in acute myeloid leukemia. Blood 138, 1830–1842. 10.1182/blood.2020009192.

54. Blöchl, C., Wang, D., Madunić, K., Lageveen-Kammeijer, G.S.M., Huber, C.G., Wuhrer, M., and Zhang, T. (2021). Integrated n-and o-glycomics of acute myeloid leukemia (Aml) cell lines. Cells 10, 3058. 10.3390/CELLS10113058/S1.

55. Blöchl, C., Wang, D., Mayboroda, O.A., Lageveen-Kammeijer, G.S.M., and Wuhrer, M. (2023). Transcriptionally imprinted glycomic signatures of acute myeloid leukemia. Cell Biosci. 13, 31-. 10.1186/S13578-023-00981-0/FIGURES/6.

56. Kikushige, Y. (2021). TIM-3 in normal and malignant hematopoiesis: Structure, function, and signaling pathways. Cancer Sci. 112, 3419–3426. 10.1111/cas.15042.

57. Tarp, M.A., Sørensen, A.L., Mandel, U., Paulsen, H., Burchell, J., Taylor-Papadimitriou, J., and Clausen, H. (2007). Identification of a novel cancer-specific immunodominant glycopeptide epitope in the MUC1 tandem repeat. Glycobiology 17, 197–209. 10.1093/GLYCOB/CWL061.

58. Greco, B., Paolella, K., Camisa, B., Malacarne, V., Falcone, L., Graziani, A., Bonini, C., Bondanza, A., and Casucci, M. (2018). Combining De-Glycosylating Agents with CAR-T Cells for Targeting Solid Tumors and Reducing Toxicity. Blood 132, 4544. 10.1182/BLOOD-2018-99-116019.

59. Liu, F.T., and Rabinovich, G.A. (2005). Galectins as modulators of tumour progression. Nat. Rev. Cancer 5, 29–41. 10.1038/NRC1527.

60. Takahashi, S. (2025). A mini-review: the role of glycosylation in acute myeloid leukemia and its potential for treatment. Oncologie 27, 689–695. 10.1515/oncologie-2025-0155.

61. Asayama, T., Tamura, H., Ishibashi, M., Kuribayashi-Hamada, Y., Onodera-Kondo, A., Okuyama, N., Yamada, A., Shimizu, M., Moriya, K., Takahashi, H., et al. (2017). Functional expression of Tim-3 on blasts and clinical impact of its ligand galectin-9 in myelodysplastic syndromes. Oncotarget 8, 88904–88917. 10.18632/ONCOTARGET.21492.

62. Wolf, Y., Anderson, A.C., and Kuchroo, V.K. (2020). TIM3 comes of age as an inhibitory receptor. Nat. Rev. Immunol. 20, 173–185. 10.1038/s41577-019-0224-6.

63. Krishnamoorthy, V., Daly, J., Kim, J., Piatnitca, L., Yuen, K.A., Kumar, B., Taherzadeh Ghahfarrokhi, M., Bui, T.Q.T., Azadi, P., Vu, L.P., et al. (2024). The glycosyltransferase ST3GAL4 drives immune evasion in acute myeloid leukemia by synthesizing ligands for the glyco-immune checkpoint receptor Siglec-9. Leuk. 2024 392 39, 346–359. 10.1038/s41375-024-02454-w.

64. Dai, Y., Cheng, Z., Pang, Y., Jiao, Y., Qian, T., Quan, L., Cui, L., Liu, Y., Si, C., Chen, J., et al. (2020). Prognostic value of the FUT family in acute myeloid leukemia. Cancer Gene Ther. 27, 70–80. 10.1038/S41417-019-0115-9.

65. Duan, C., Fukuda, T., Isaji, T., Qi, F., Yang, J., Wang, Y., Takahashi, S., and Gu, J. (2020). Deficiency of core fucosylation activates cellular signaling dependent on FLT3 expression in a Ba/F3 cell system. FASEB J. 34, 3239–3252. 10.1096/FJ.201902313RR.

66. Liu, J., and Gu, J. (2024). Importance of PTM of FLT3 in acute myeloid leukemia: Importance of PTM of FLT3 in acute myeloid leukemia. Acta Biochim. Biophys. Sin. (Shanghai). 56, 1199. 10.3724/ABBS.2024112.

67. Schauner, R., Cress, J., Hong, C., Wald, D., and Ramakrishnan, P. (2024). Single cell and bulk RNA expression analyses identify enhanced hexosamine biosynthetic pathway and O-GlcNAcylation in acute myeloid leukemia blasts and stem cells. Front. Immunol. 15, 1327405. 10.3389/FIMMU.2024.1327405/BIBTEX.

68. Asthana, A., Ramakrishnan, P., Vicioso, Y., Zhang, K., and Parameswaran, R. (2018). Hexosamine Biosynthetic Pathway Inhibition Leads to AML Cell Differentiation and Cell Death. Mol. Cancer Ther. 17, 2226–2237. 10.1158/1535-7163.MCT-18-0426.

69. Park, S., Ho, C.E., Darnell, E.P., Wolff, A.N., Takei, H., Birocchi, F., Bouffard, A.A., Salas-Benito, D., Escobar, G., Leick, M.B., et al. (2025). Tuning CAR-T cells by targeting cancer-associated glycan in pancreatic cancer. Nat. Commun. 2025 161 16, 11246-. 10.1038/s41467-025-66102-2.

70. Molica, M., Perrone, S., Mazzone, C., Niscola, P., Cesini, L., Abruzzese, E., and de Fabritiis, P. (2021). CD33 Expression and Gentuzumab Ozogamicin in Acute Myeloid Leukemia: Two Sides of the Same Coin. Cancers (Basel). 13. 10.3390/CANCERS13133214.

71. Dutour, A., Marin, V., Pizzitola, I., Valsesia-Wittmann, S., Lee, D., Yvon, E., Finney, H., Lawson, A., Brenner, M., Biondi, A., et al. (2012). In vitro and in vivo antitumor effect of anti-CD33 chimeric receptor-expressing EBV-CTL against CD 33 + acute myeloid leukemia. Adv. Hematol. 2012. 10.1155/2012/683065.

72. Marin, V., Pizzitola, I., Agostoni, V., Attianese, G.M.P.G., Finney, H., Lawson, A., Pule, M., Rousseau, R., Biondi, A., and Biagi, E. (2010). Cytokine-induced killer cells for cell therapy of acute myeloid leukemia: improvement of their immune activity by expression of CD33-specific chimeric receptors. Haematologica 95, 2144–2152. 10.3324/HAEMATOL.2010.026310.

73. Katsarou, A., Sjöstrand, M., Naik, J., Mansilla-Soto, J., Kefala, D., Kladis, G., Nianias, A., Ruiter, R., Poels, R., Sarkar, I., et al. (2021). Combining a CAR and a chimeric costimulatory receptor enhances T cell sensitivity to low antigen density and promotes persistence. Sci. Transl. Med. 13. 10.1126/SCITRANSLMED.ABH1962.

74. Muliaditan, T., Halim, L., Whilding, L.M., Draper, B., Achkova, D.Y., Kausar, F., Glover, M., Bechman, N., Arulappu, A., Sanchez, J., et al. (2021). Synergistic T cell signaling by 41BB and CD28 is optimally achieved by membrane proximal positioning within parallel chimeric antigen receptors. Cell reports. Med. 2. 10.1016/J.XCRM.2021.100457.

75. Drent, E., Poels, R., Ruiter, R., Van De Donk, N.W.C.J., Zweegman, S., Yuan, H., De Bruijn, J., Sadelain, M., Lokhorst, H.M., Groen, R.W.J., et al. (2019). Combined CD28 and 4-1BB Costimulation Potentiates Affinity-tuned Chimeric Antigen Receptor-engineered T Cells. Clin. Cancer Res. 25, 4014–4025. 10.1158/1078-0432.CCR-18-2559.

76. Kloss, C.C., Condomines, M., Cartellieri, M., Bachmann, M., and Sadelain, M. (2013). Combinatorial antigen recognition with balanced signaling promotes selective tumor eradication by engineered T cells. Nat. Biotechnol. 31, 71–75. 10.1038/nbt.2459.

77. Grada, Z., Hegde, M., Byrd, T., Shaffer, D.R., Ghazi, A., Brawley, V.S., Corder, A., Schönfeld, K., Koch, J., Dotti, G., et al. (2013). TanCAR: A novel bispecific chimeric antigen receptor for cancer immunotherapy. Mol. Ther. - Nucleic Acids 2. 10.1038/mtna.2013.32.

78. Perriello, V.M., Rotiroti, M.C., Pisani, I., Galimberti, S., Alberti, G., Pianigiani, G., Ciaurro, V., Marra, A., Sabino, M., Tini, V., et al. (2023). IL-3-zetakine combined with a CD33 costimulatory receptor as a dual CAR approach for safer and selective targeting of AML. Blood Adv. 7, 2855–2871. 10.1182/BLOODADVANCES.2022008762.

