## Supplemental File for "Differential TIM-3 glycosylation enables specific dual targeting CAR-T therapy in acute myeloid leukemia"

#### Supplemental Information

**Supplemental Figures and Legends**

### Figure S1

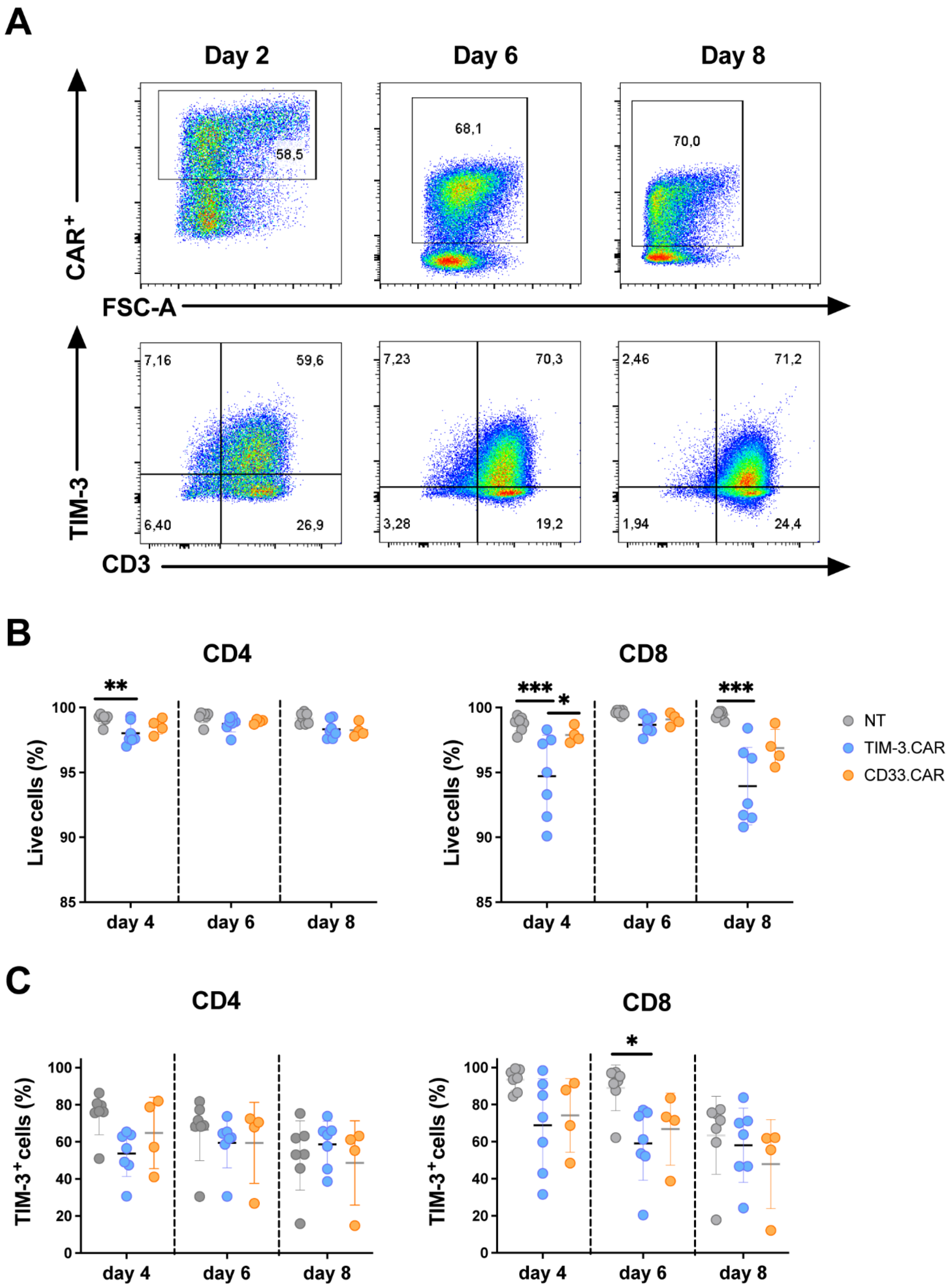

**Figure S1.** ***In vitro*** **fratricide assessment of TIM-3.CAR-CIK cells on CD4$^+$ and CD8$^+$ cell subsets, related to Figure 1.**

(A) Representative flow cytometry plots of CD3$^+$ CAR$^+$ and TIM-3$^+$ expression on TIM-3.CAR-CIK cells over the first days after CAR transduction.
(B) Viability of TIM-3.CAR-CIK cells compared to CD33.CAR-CIK and NT cells at days 4,6 and 8 post-transduction on CD4$^+$ and CD8$^+$ T cell subsets.
(C) Expression of TIM-3 in TIM3.CAR-CIK, CD33.CAR-CIK and NT cells over the first 8 days of CAR-CIK differentiation on CD4$^+$ and CD8$^+$ T cell subsets (n=7 for TIM3.CAR-CIK and NT cells, n=4 for CD33.CAR-CIK cells).

Data are shown as individual values ± SD. Statistical significance was assessed using repeated-measured two-way ANOVA with Bonferroni's post hoc test. *p = 0.01, **p < 0.001, ***p = 0.0001.

### Figure S2

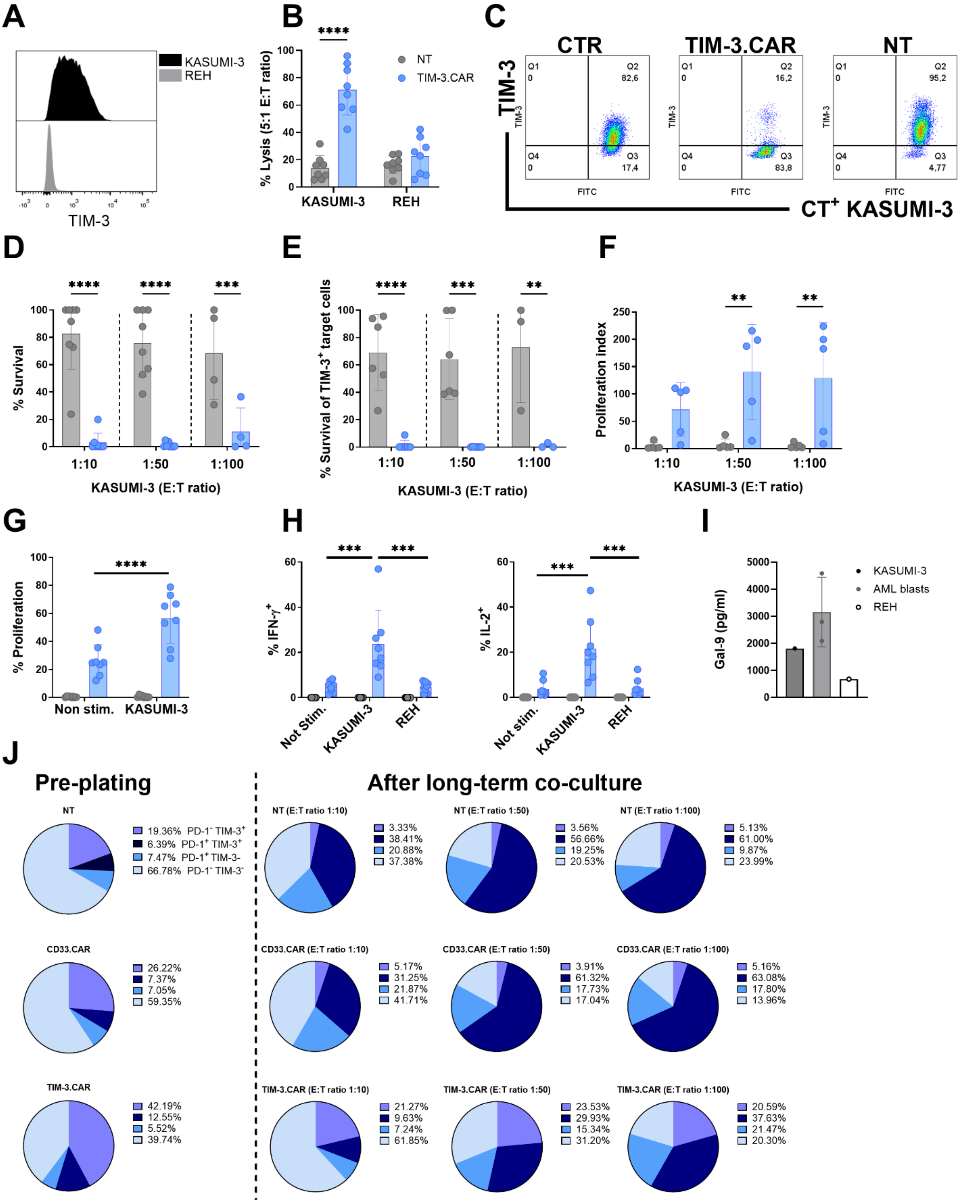


**Figure S2. *In vitro* antileukemic activity of TIM-3.CAR-CIK cells against TIM-3⁺ KASUMI-3 cell line, related to Figure 2.**

(A) TIM-3 expression on KASUMI-3 cell line assessed by flow cytometry. REH, cell line served as negative control.

(B) Short-term killing assay of TIM-3.CAR-CIK cells against KASUMI-3 or control REH cells compared with NT cells. Target cell lysis was assessed by flow cytometry (E:T 5:1, n = 8)

(C) Representative flow cytometry plots showing the residual percentage of TIM-3+ KASUMI-3 cells cultured alone or after 4 hours co-culture with TIM-3.CAR-CIK or NT cells.
(D) Long-term killing assay of TIM-3.CAR-CIK cells against KASUMI-3 cells compared with NT cells. KASUMI-3 survival was evaluated by flow cytometry (ET 1:10 and 1:50, n = 8; E:T 1:100, n = 4).
(E) Survival of TIM-3+ KASUMI-3 cells after 7-day co-culture with TIM-3.CAR-CIK or NT cells (E:T 1:10 and 1:50, n = 6; E:T 1:100, n = 3).
(F) Proliferation index of TIM-3.CAR-CIK cells compared with NT cells after 7-day co-culture with KASUMI-3 cells. Proliferation was quantified by flow cytometry as the ratio of CD3+ cells recovered from experimental wells to those from CIK-only control wells (ET 1:10, 1:50 and 1:100, n = 5).
(G) Proliferation of TIM-3.CAR-CIK cells assessed by Ki67 staining after 72 hours co-culture with KASUMI-3 (E:T 1:1, n = 8).
(H) Cytokine production (IFN-γ, IL-2) after 5 hours co-culture of TIM-3.CAR-CIK or NT cells with primary AML blasts (E:T 1:3, n = 8).
(I) Concentration of Gal-9 produced by KASUMI-3, primary AML blasts (n =3) and REH cells, measured by ELISA.
(J) Pie charts representing the expression of PD-1 and TIM-3 on NT and TIM3.CAR-CIK cells before plating the long-term killing assay and after 7 days co-culture with KASUMI-3 cells at all three E:T ratios tested.

Data are shown as individual values ± SD. Statistical significance was assessed using repeated-measured two-way ANOVA with Bonferroni's post hoc test. ** $p<0.001$, *** $p=0.0001$ and **** $p<0.0001$.

### Figure S3

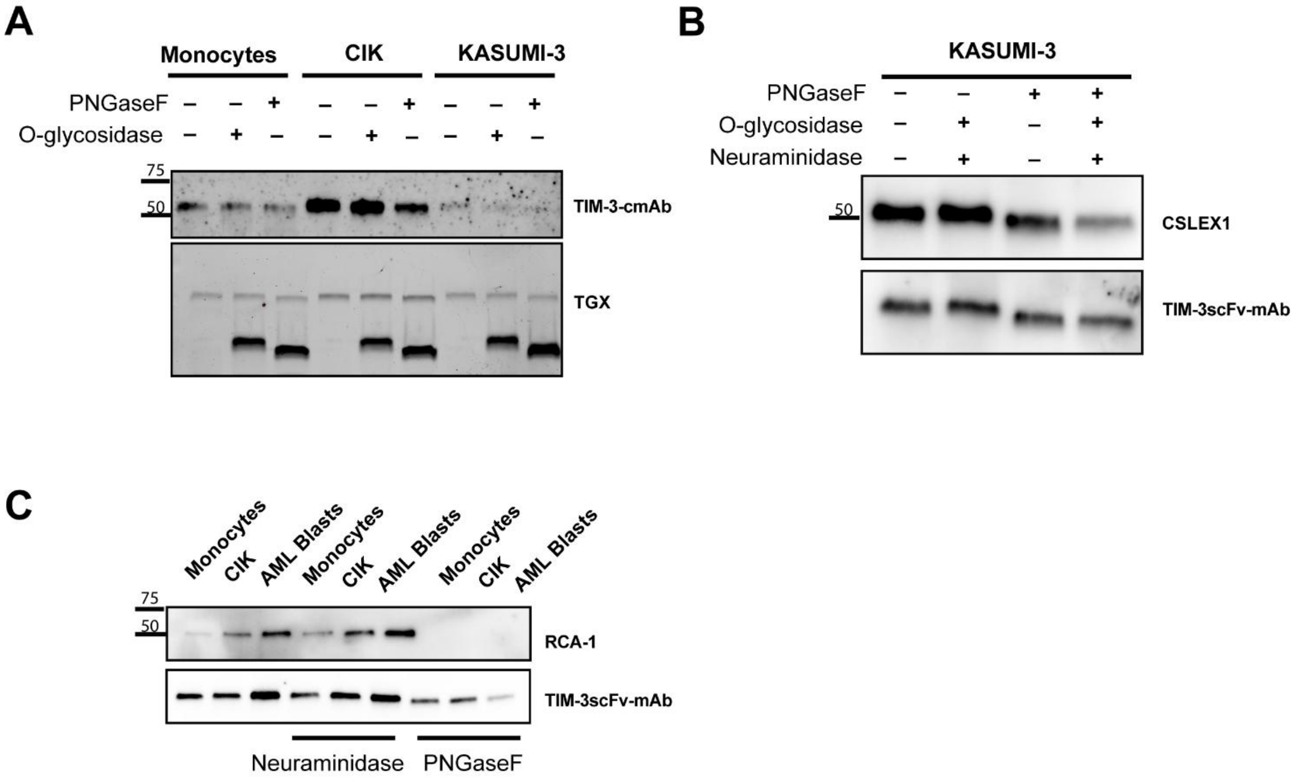


**Figure S3. Additional lectin/antibody probing of TIM-3 glycoforms and terminal galactose exposure, related to Figures 3 and 4**.

(A) Immunoblot detection of TIM-3 species in monocytes, CIK cells, and KASUMI-3 cells using TIM-3-cmAb following PNGase F or *O*-glycosidase treatment. TGX stain-free total protein signal is shown as a loading/normalization control.

(B) TIM-3 immunoprecipitates from KASUMI-3 cells treated with PNGase F, *O*-glycosidase, and/or neuraminidase and probed with CSLEX1 (sialyl-Lewis X) and TIM-3scFv-mAb.

(C) RCA-1 lectin probing of TIM-3 material from monocytes, CIK cells, and KASUMI-3 cells following neuraminidase or PNGase F treatment, with TIM-3scFv-mAb shown as a control readout.

Immunoblot and lectin blot experiments (A-C) were repeated in three independent biological replicates with similar results.

### Figure S4

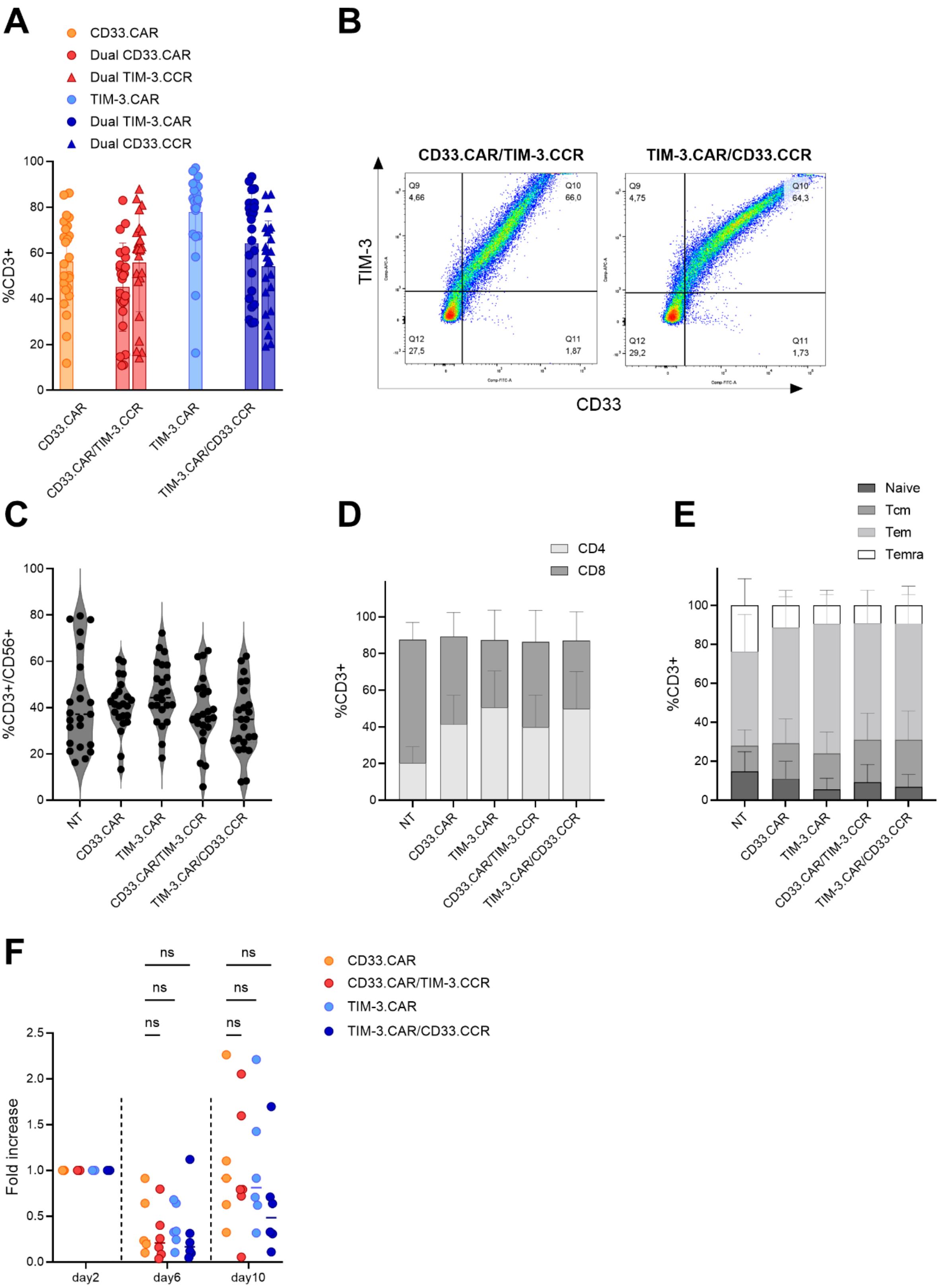
A
CD33.CAR
Dual CD33.CAR
Dual TIM-3.CCR
TIM-3.CAR
Dual TIM-3.CAR
Dual CD33.CCR
%CD3+
CD33.CAR
CD33.CAR/TIM-3.CCR
TIM-3.CAR
TIM-3.CAR/CD33.CCR
B
CD33.CAR/TIM-3.CCR
TIM-3.CAR/CD33.CCR
TIM-3
CD33
Q9 4,66
Q10 66,0
Q12 27,5
Q11 1,87
Q9 4,75
Q10 64,3
Q12 29,2
Q11 1,73
C
%CD3+/CD56+
NT
CD33.CAR
TIM-3.CAR
CD33.CAR/TIM-3.CCR
TIM-3.CAR/CD33.CCR
D
CD4
CD8
%CD3+
E
Naive
Tcm
Tem
Temra
%CD3+
F
ns
Fold increase
day2
day6
day10
CD33.CAR
CD33.CAR/TIM-3.CCR
TIM-3.CAR
TIM-3.CAR/CD33.CCR

**Figure S4. Phenotypic characterization of IF-BETTER dual targeting CD33.CAR/TIM-3.CCR and TIM-3.CAR/CD33.CCR constructs, related to Figure 6.**

(A) Summary of CD33.CAR/TIM-3.CCR and TIM-3.CAR/CD33.CCR expression percentage, compared with CD33.CAR and TIM-3.CAR, used as internal controls, at the end of the culture (n = 18).
(B) Flow cytometry plots representative of CAR and CCR co-expression in both Dual CAR-CIK cells at the end of culture (n = 18).
(C) Violin plots showing the percentage of $CD3^{+}$ $CD56^{+}$ cells in Dual CAR-CIK cells compared with CD33.CAR-CIK and TIM-3.CAR-CIK cells, used as controls (n = 18).
(D) Relative percentages of $CD4^{+}$ and $CD8^{+}$ populations in all CAR-CIK cells (n = 18).
(E) Representation of all CAR-CIK cell memory phenotype, divided in $T_{naive}$, $T_{central\ memory}$, $T_{effector\ memory}$ and $T_{EMRA}$ populations (n = 18).
(F) Fold increase of Dual CAR-CIK cells compared to CD33.CAR and TIM-3.CAR-CIK cells at days 2, 6, and 10 of the culture (n = 6).

Data are shown as individual values ± SD. Statistical significance was assessed using repeated-measured two-way ANOVA with Bonferroni’s post hoc test. ns, not significant.

### Figure S5

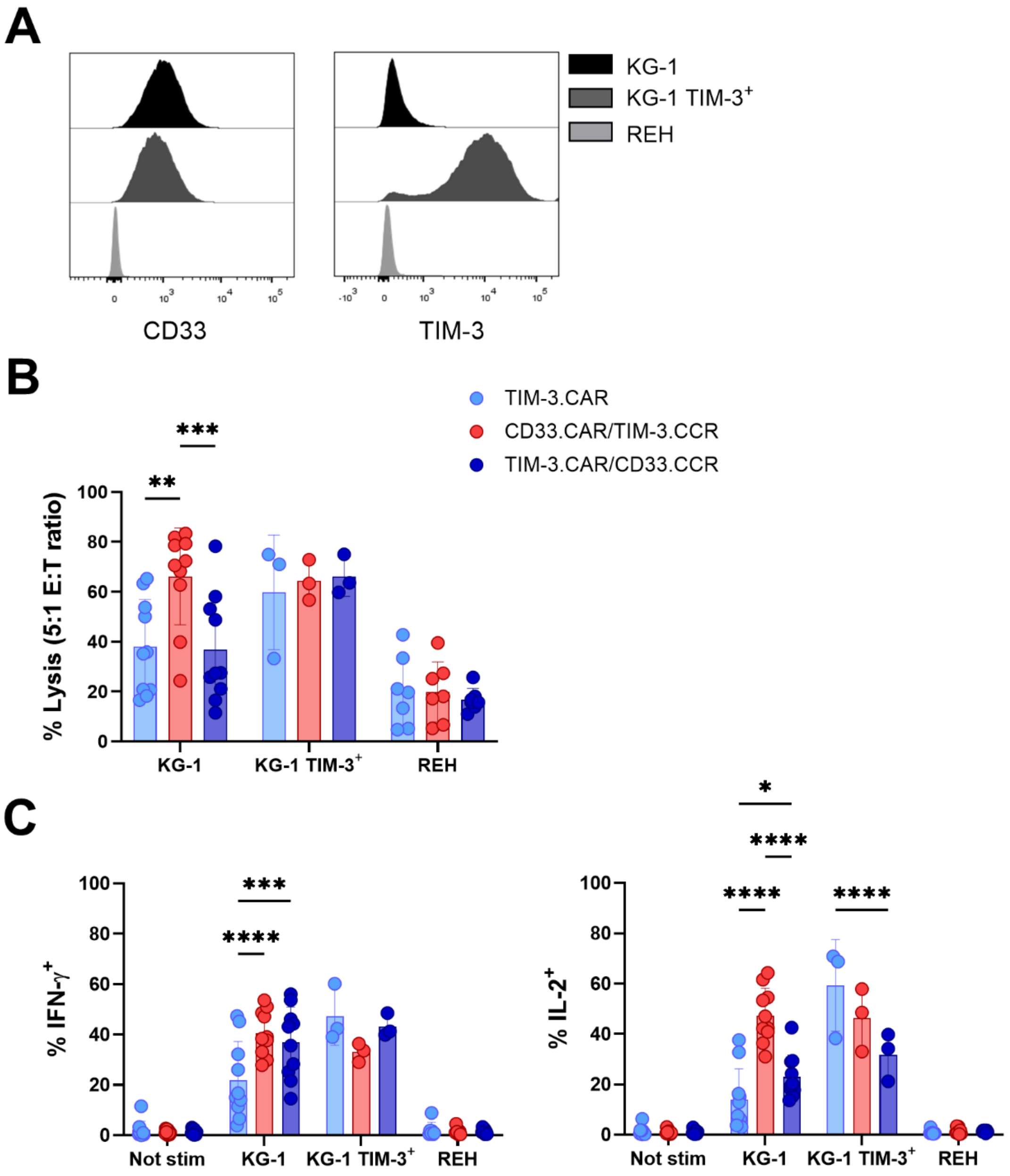


**Figure S5.** ***In vitro*** **antileukemic activity of Dual CAR-CIK cells against KG-1 and KG-1 TIM-3**$^{+}$ **cell lines, related to Figure 6.**

(A) CD33 and TIM-3 expression on KG-1 and KG-1 TIM-3$^{+}$ cell lines assessed by flow cytometry. REH, cell line served as negative control.

(B) Short-term killing assay of TIM-3.CAR-CIK cells and Dual CAR-CIK cells against target AML cell lines or control REH cells. Target cell lysis was assessed by flow cytometry (E:T 5:1, n = 10 for KG-1 cells, n = 3 for KG-1 TIM-3$^{+}$ cells and n = 7 for REH cells)

(C) Cytokine production (IFN-γ, IL-2) after 5 hours co-culture of TIM-3.CAR-CIK or Dual CAR-CIK cells with KG-1, KG-1 TIM-3$^{+}$ or REH cell lines (E:T 1:3, n = 10 for KG-1 cells, n = 3 for KG-1 TIM-3$^{+}$ cells and n = 7 for REH cells).

Data are shown as individual values ± SD. Statistical significance was assessed using repeated-measured two-way ANOVA with Bonferroni's post hoc test. *p = 0.01, **p < 0.001, ***p = 0.0001 and ****p < 0.0001.

#### Figure S6

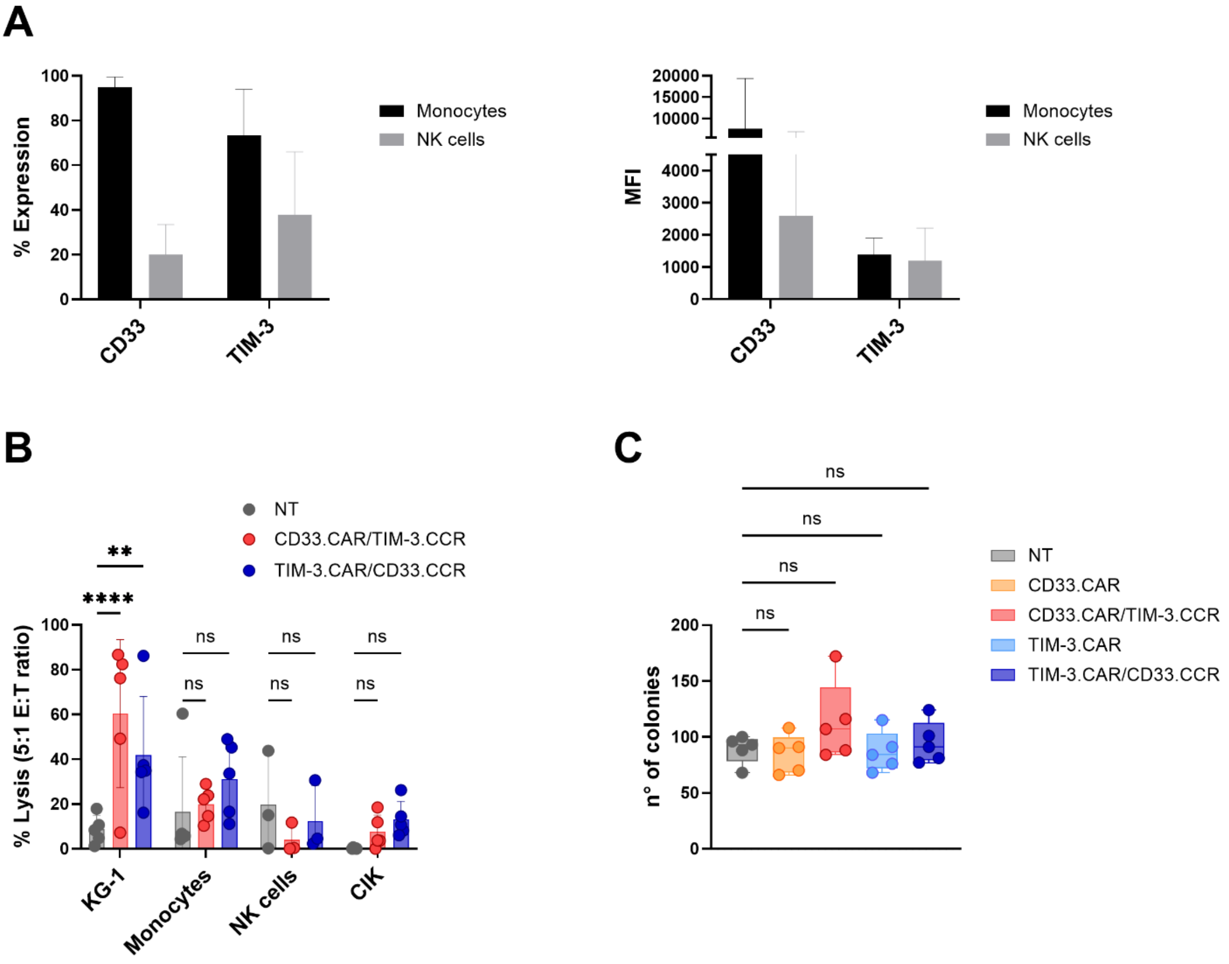


**Figure S6. Minimal *in vitro* cytotoxicity of Dual CAR-CIK cells toward CD33⁺ TIM-3⁺ healthy immune cells and HSCs, related to Figure 3.**

(A) Frequency (left) and MFI (right) of CD33 and TIM-3 expression on healthy immune subsets (monocytes and NK cells) assessed by flow cytometry.
(B) Short-term killing assay of Dual CAR-CIK cells against monocytes, NK- or CIK cells compared with NT cells. KG-1 cell line served as positive control. Target cell lysis was evaluated by flow cytometry (E:T 5:1, n = 5 for KG-1 cells, monocytes and CIK cells, n = 3 for NK cells).
(C) Colony Forming Unit (CFU) assay of all CAR-CIK cells compared with NT cells. Total number of colonies was assessed by microscopy.

Data are shown as individual values ± SD. Statistical significance was assessed using repeated-measured two-way ANOVA with Bonferroni's post hoc test. **p < 0.001 and ****p < 0.0001.
